# Toxic metal, Lead (Pb) alters columella-specific auxin distribution inhibiting root growth and gravitropic response in Arabidopsis and maize

**DOI:** 10.1101/2025.05.09.653208

**Authors:** Julia Zheku, Scottie Mello, Olivia S. Hazelwood, Grace Delpit, Erin E. Sparks, Norman B. Best, M. Arif Ashraf

## Abstract

Toxic metal contamination is pervasive throughout global soils and of significant concern due to its high abundance in agricultural lands. Contaminated soil detrimentally impacts the crops that grow in them and the humans that consume these crops. There is a need for engineering of plants able to grow in the presence of toxic metals while sequestering these contaminants away from edible tissues. However, engineering efforts require a detailed mechanistic understanding of plant responses to these toxins, which are currently lacking. In this study, we discovered that there are dissimilar root growth responses to different toxic metals. We found that the toxic metal lead (Pb) targets conserved cellular and developmental processes in evolutionary diverse plant systems - the model plant *Arabidopsis thaliana* and crop plant *Zea mays*. In *Arabidopsis thaliana*, Pb inhibits auxin transporter PIN FORMED 3 (PIN3) protein trafficking in the columella cells, which in turn alters the phytohormone auxin distribution, and consequently inhibits cell cycle progression and disrupts root gravitropic responses. Both root growth and gravitropic responses are critical for soil exploration, which requires plants to live and thrive in harsh environments.

## Introduction

Metal-contaminated soils cause harm to the plants that grow in them and the humans that consume these plants as food. Most harmful among these metal contaminants are toxic metals like chromium (Cr), cadmium (Cd), lead (Pb), cesium (Cs), and arsenic (As) (Hou et al. 2020, 2025). A recent study reported that 14 to 17% of agricultural land is contaminated by toxic metals across the globe and this contamination affects one third of the world’s population (Hou et al. 2025). These contaminants are particularly challenging to mitigate since they share a chemical similarity with essential minerals, thus limiting the ability of plants to distinguish elemental friend and foe (Ashraf et al. 2020, 2021). For instance, As utilizes phosphate transporters (Catarecha et al. 2007) and Cs takes advantage of potassium transporters (Adams et al. 2019) due to their chemical similarity. The iron and manganese transporter, IRON-REGULATED TRANSPORTER1 (IRT1), can also transport Cd and Pb due to its broad substrate non-specificity (Korshunova et al. 1999; Rogers et al. 2000; Cointry and Vert 2019). Despite the negative impacts on plants and humans, there is little known about how plants respond to toxic metal stress and whether the responses are shared or divergent among different metals and different plant species. Uncovering these mechanisms is critical for future mitigation efforts.

Because of its close contact with the water and soil, the root is the first organ to get exposed to toxic metals. Toxic metals, such as Cd, Pb, As, Cr, and Cs all inhibit the root growth of *Arabidopsis thaliana* (Arabidopsis) (Jiang and Liu 2000; Fukao et al. 2011; Yuan and Huang 2016a; Song et al. 2017; Ashraf et al. 2021; Ortiz-Luevano et al. 2021; Zheng et al. 2021; Tao et al. 2025; Thiébaut et al. 2025). Mechanistically, a reduction in root growth can be due to reduced cell division and/or reduced cell elongation (Rahman et al. 2007; Ashraf and Rahman 2019; Ashraf et al. 2021). Some studies reported that toxic metal-mediated root growth inhibition happens due to the slower cell division activity in the meristematic region of the Arabidopsis and maize root (Jiang and Liu 2000; Song et al. 2017; Ashraf et al. 2021; Ortiz-Luevano et al. 2021; Zheng et al. 2021; Tao et al. 2025; Thiébaut et al. 2025). Among these reports, a portion of studies used pCYCB1;1-GUS/GFP as molecular reporter to investigate cell cycle regulation (Yuan and Huang 2016b; Song et al. 2017; Ashraf et al. 2021; Zheng et al. 2021; Tao et al. 2025). Furthermore, a handful of studies took a careful look at the cell elongation of the epidermal cells after toxic metal exposure (Fukao et al. 2011; Ashraf et al. 2021; Tao et al. 2025). Although these studies provided an overall idea about the cell cycle regulation and differentiation in root tissue during toxic metal stress, a comprehensive cell cycle profiling along with cell elongation responses has not been reported. The recent generation of cell cycle reporter lines, such as Cytrap (differentiating S/G2 and G2/M phases), PCNA1-sGFP (G1/G2, early and late S phases), and PlaCCI (<u>pla</u>nt <u>c</u>ell <u>c</u>ycle indicator) can help identify the exact points that are being effected in cell cycle regulation during toxic metal stress (Yin et al. 2014; Yokoyama et al. 2016; Desvoyes et al. 2020; Hazelwood and Ashraf 2024).

Upon exposure to toxic metals, the differential regulation of phytohormones, such as auxin (Yuan et al. 2013; Li et al. 2015; Wang et al. 2015; Ashraf et al. 2020), abscisic acid (ABA) (Monni et al. 2001; Atici et al. 2005; Fediuc et al. 2005; Kim et al. 2014; Wang et al. 2014), and ethylene (Schellingen et al. 2014; Khan et al. 2015), has been reported in a wide range of plant species. This suggests a conservation of toxic metal-induced phytohormone responses similar to the conserved role of phytohormone-mediated adaptation to the environment during evolution (Arabia and Munné-Bosch 2026). Phytohormones are often synthesized in one part of the plant and moved throughout the plant via transporters, which are rapidly recycled and relocalized to redirect hormone flux and alter downstream gene expression. Changes in phytohormones may be regulated at the biosynthesis, transport, or signaling level.

Among the most well characterized phytohormones is auxin. Auxin biosynthesis starts with the production of Tryptophan, originating with the enzyme complex anthranilate synthase converting chorismate to anthranilate, which is the rate-limiting step in tryptophan biosynthesis (Mano and Nemoto 2012). In Arabidopsis, WEAK ETHYLENE INSENSITIVE 2 (WEI2) and WEI7 encode the alpha- and beta-subunits of this synthase and thus mutation of these genes disrupts auxin production. After the production of Tryptophan, the enzyme TRYPTOPHAN AMINOTRANSFERASE OF ARABIDOPSIS 1 (TAA1) converts Tryptophan to indole-3-pyruvate, the first step towards the production of auxin. In Arabidopsis, the WEI8 gene encodes for TAA1, and mutation of this gene disrupts auxin production (Stepanova et al. 2008). There have been other pathways proposed to synthesize auxin in plants, but this is the main pathway in Arabidopsis (Mashiguchi et al. 2011). However, disruption of auxin biosynthesis causes pleiotropic effects, making it difficult to distinguish its role in specific biological responses.

After biosynthesis, auxin is transported across cells through a series of influx and efflux transporters. Influx transporters include AUXIN1/LIKE AUX1 (AUX1/LAX) transporters. In Arabidopsis, there are four AUX1/LAXs with two having specific functions in the root (AUX1 and LAX3) (Matthes et al. 2019a). Efflux transporters include PIN-FORMED (PIN) transporters. In Arabidopsis, there are eight PINs, which are divided into canonical (PIN1, EIR1/PIN2, PIN3, PIN4 and PIN7) and non-canonical (PIN5, PIN6 and PIN8) based on their sequence and localization (Mravec et al. 2009; Ding et al. 2012; Ditengou et al. 2018; Matthes et al. 2019b; Ung et al. 2022). Canonical PINs are plasma membrane localized and control movement between cells, whereas non-canonical PINs are internally localized. Auxin transporters are continuously recycled from the plasma membrane to enable rapid responses and redirect auxin flux. One approach to determine the role of transporters in a biological response is to disrupt membrane trafficking by treatment with Brefeldin A (BFA) (Geldner et al. 2001).

Nuclear auxin signaling relies on a de-repression mechanism. In the absence of auxin, Aux/IAA repressors bind to AUXIN RESPONSE FACTOR (ARF) transcription factors preventing any downstream responses. Once inside the cell, auxin binds with the TRANSPORT INHIBITOR RESPONSE 1/AUXIN SIGNALING F-BOX (TIR1/AFB) receptor (Kepinski and Leyser 2005). This binding triggers the ubiquitination and subsequent degradation of the Aux/IAA repressor, freeing the ARF transcription factors to modulate gene expression. A key component of the degradation pathway in Arabidopsis is the AUXIN-RESISTANT 1 (AXR1) gene (Leyser et al. 1993; Lavy and Estelle 2016). Disruption of either the receptor or the degradation pathway results in a disruption of auxin downstream signaling. We have a detailed understanding of the auxin pathway in Arabidopsis, but the characterization in other plants such as the crop *Zea mays* (maize) is less well defined.

In this study, we tested the root growth and phytohormone response of Arabidopsis and maize after exposure to four major toxic metals (Cd, Pb, As, and Cr), and identified similar but distinct responses. Among these toxic metals, Pb caused distinct root-specific phenotypes – primary root growth inhibition and altered gravitropic response, which were attributed to increased auxin accumulation in both species. Leveraging the molecular genetic tools in Arabidopsis, we showed that Pb specifically inhibits the G0/G1 phase of the cell cycle to reduce root growth without altering cell elongation. Despite the increased auxin accumulation in response to Pb, we showed that Pb altered auxin distribution in the root, which causes aberrant gravitropic responses. Reverse genetic screening of auxin signaling, biosynthesis, and transport mutant response to Pb-exposure, identified PIN3 as a major target for Pb-mediated root growth responses. Furthermore, we showed that Pb inhibited PIN3 trafficking in the gravi-sensing columella cells and consequently altered the auxin response required for maintaining gravitropic response. The negative impact of Pb on primary root growth and gravitropic response results in the inability of roots to successfully navigate the soil environment, which impairs the ability of plants to survive in the presence of Pb. Altogether, this study highlights a broadly conserved response to Pb in evolutionary diverse plant organisms and defines a foundational role for PIN3-mediated auxin distribution in these responses.

## Results

### Plant roots have differential growth responses to toxic metals, which is shared among plant species

To evaluate the impact of toxic metals on two divergent plant species, Arabidopsis and maize plants were grown in the presence of four different metals, As, Cd, Cr, and Pb and As, at increasing dose concentrations (0, 10, 100, 1000 µM for Pb; 0, 1, 10, 100 µM for As, Cd and Cr) and evaluated at 24h, 28h, and 72h after exposure. For As, both species showed reduced root growth, although Arabidopsis showed greater sensitivity with earlier responses at lower concentrations compared to maize (Supplemental Figure S1). For Cd, both species showed reduced root growth, although maize showed greater sensitivity with earlier responses at lower concentrations compared to Arabidopsis (Supplemental Figure S2). For Cr, Arabidopsis showed dramatic reductions in root growth early and at low concentrations, whereas maize showed a minimal root growth reduction response to this toxic metal even at high concentrations (Supplemental Figure S3). For Pb, 100µM Pb inhibited primary root growth in both Arabidopsis and maize at 24h, 48h, and 72h time points (Figure 1). However, maize was more sensitive to Pb, as seen by the reduction in root growth upon exposure to 10µM Pb, which was not observed in Arabidopsis (Figure 1).

**Figure 1.**
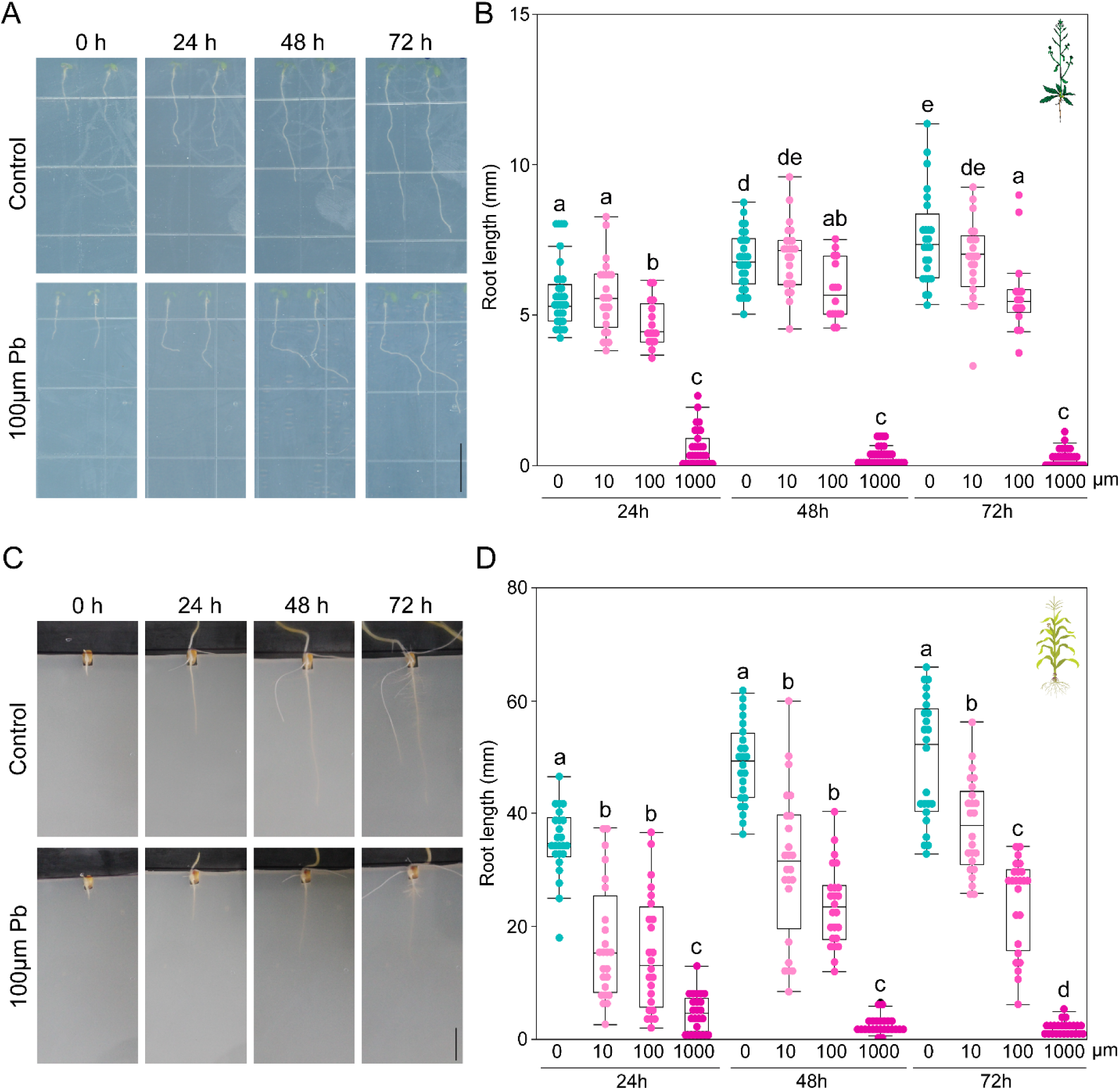
Lead inhibits root growth in a dose- and time-dependent manner. (A) 3-days-old Arabidopsis seedlings were transferred to control and 100µM Pb plates to observe the primary root growth at 24h, 48h, and 72h time points. Scale bar = 10mm. (B) Quantification of primary root growth of Arabidopsis after incubating at control, 10µM Pb, 100µM Pb, and 1000µM Pb conditions for 24h, 48h, and 72h time points. Statistical test is performed based on Tukey’s Honest test. Groups labeled with the same letter are not statistically different from each other (alpha = 0.05). Boxplots show median values (center line), 25th to 75th interquartile range (box) and 1.5*interquartile range (whiskers). (C) Primary root growth of maize at 24h, 48h, and 72h time points in control and 100µM Pb conditions. Scale bar = 20mm. (D) Quantification of primary root growth of maize after incubating at control, 10µM Pb, 100µM Pb, and 1000µM Pb conditions for 24h, 48h, and 72h time points. Statistical test is performed based on Tukey’s Honest test. Groups labeled with the same letter are not statistically different from each other (alpha = 0.05). Boxplots show median values (center line), 25th to 75th interquartile range (box) and 1.5*interquartile range (whiskers).

### Lead increases auxin levels in the primary root of Arabidopsis and maize

Root growth data suggests that maize responses to As, Cr, and Cd are distinct from Pb, with maize showing a reduced tolerance to Pb. We hypothesized that these differences would be reflected in the root phytohormone profiles after exposure (48h) to these toxic metals (10µM As, 10µM Cr, 25µM Cd, and 100µM Pb). Indeed, maize roots exposed to As have significantly elevated 12-hydroxyjasmonic acid (12OHJA), 1-aminocyclopropane-1-carboxylic acid (1-ACC), indole-3-acetic acid (IAA), and salicylic acid (SA), but significantly reduced gibberellin12 (GA12) (Table 1). Whereas Cd caused elevated jasmonic acid (JA), 1-ACC, but reduced abscisic acid (ABA), GA53, GA12, and SA; Cr caused elevated 1-ACC, and reduced ABA, GA53, and GA12 (Table 1). Lastly, Pb increased JA, IAA, and SA, but reduced GA12 (Table 1, Supplemental Figure 4). We also measured the sterol and brassinosteroid intermediates isofucosterol, stigmasterol, 24-methylenecholesterol, campesterol, campestanol, and 6-oxocampestanol in maize roots, but these were unchanged when treated with any toxic metal (Supplemental Table 1)

**Table 1:** Hormone quantification from maize root during control, As, Cd, Cr, and Pb conditions. Higher than control and lower than control is highlighted with cyan and magenta, respectively.

| Content<br>(ng/g<br>FW) | *Zea mays* |  |  |  |  |
| --- | --- | --- | --- | --- | --- |
|  | Control | As | Cd | Cr | Pb |
| **12OHJA** | 2921.5 ±<br>1319.39 | 6918.43 ±<br>863.85 (*) | 1338.84 ±<br>432.87 | 308.32 ±<br>161.45 | 5382.48 ±<br>780.47 |
| **JA** | 650.05 ±<br>133.07 | 1187.59 ±<br>288.53 | 1243.64 ±<br>184.64 (*) | 784.66 ±<br>94.26 | 1620.92 ±<br>151.14 (**) |
| **1-ACC** | 101.85 ±<br>29.04 | 290.75 ±<br>42.37 (**) | 362.099 ±<br>45.51 (**) | 312.88 ±<br>66.54 (*) | 248.78 ±<br>88.29 |
| **ABA** | 214.21 ±<br>50.78 | 154.74 ±<br>33.79 | 31.24 ±<br>5.22 (*) | 23.33 ±<br>10.12 (*) | 150.5 ±<br>32.69 |
| **GA53** | 68.83 ±<br>19.13 | 68.17 ±<br>8.41 | 24.49 ±<br>7.23 (*) | 19.40 ±<br>6.96 (*) | 68.45 ±<br>25.55 |
| **GA12** | 1344.9 ±<br>265.02 | 404.21 ±<br>115.29 (*) | 64.41 ±<br>12.82 (*) | 24.58 ±<br>11.54 (*) | 741.48 ±<br>122.11 (*) |
| **IAA** | 27.26 ±<br>9.51 | 66.77 ±<br>13.89 (*) | 19.37 ±<br>7.83 | 19.52 ±<br>2.05 | 218.84 ±<br>52.78 (*) |
| **SA** | 152.22 ±<br>23.51 | 899.71 ±<br>134.43 (**) | 31.30 ±<br>17.43 (**) | 199.84 ±<br>63.91 | 983.38 ±<br>178.79 (*) |

Auxin (also known as IAA) is known to be involved in toxic metal responses such as inhibited root growth (Ashraf et al. 2020; Tao et al. 2025). Indeed, the highest levels of IAA were identified in response to Pb, which is consistent with the most severe growth reduction (Table 1). To determine if the phytohormone responses to Pb were consistent in Arabidopsis, phytohormone profiles were quantified from Arabidopsis roots after 48h exposure to lead as compared to control treatment(Table 2). Similar to maize, there was a significant increase in JA, and IAA after Pb exposure (Table 2). In contrast to maize, there was also an increase in 1-ACC and no changes in other phytohormones (Table 2). IAA levels increased ∼8-fold in maize and ∼4-fold in Arabidopsis in response to Pb (Figures 2A and 2B). Given the well-document role of auxin in root development in both Arabidopsis (Roychoudhry and Kepinski 2022) and maize (Cowling et al. 2023); we hypothesized that auxin is a key mediator of the Pb-response.

**Figure 2.**
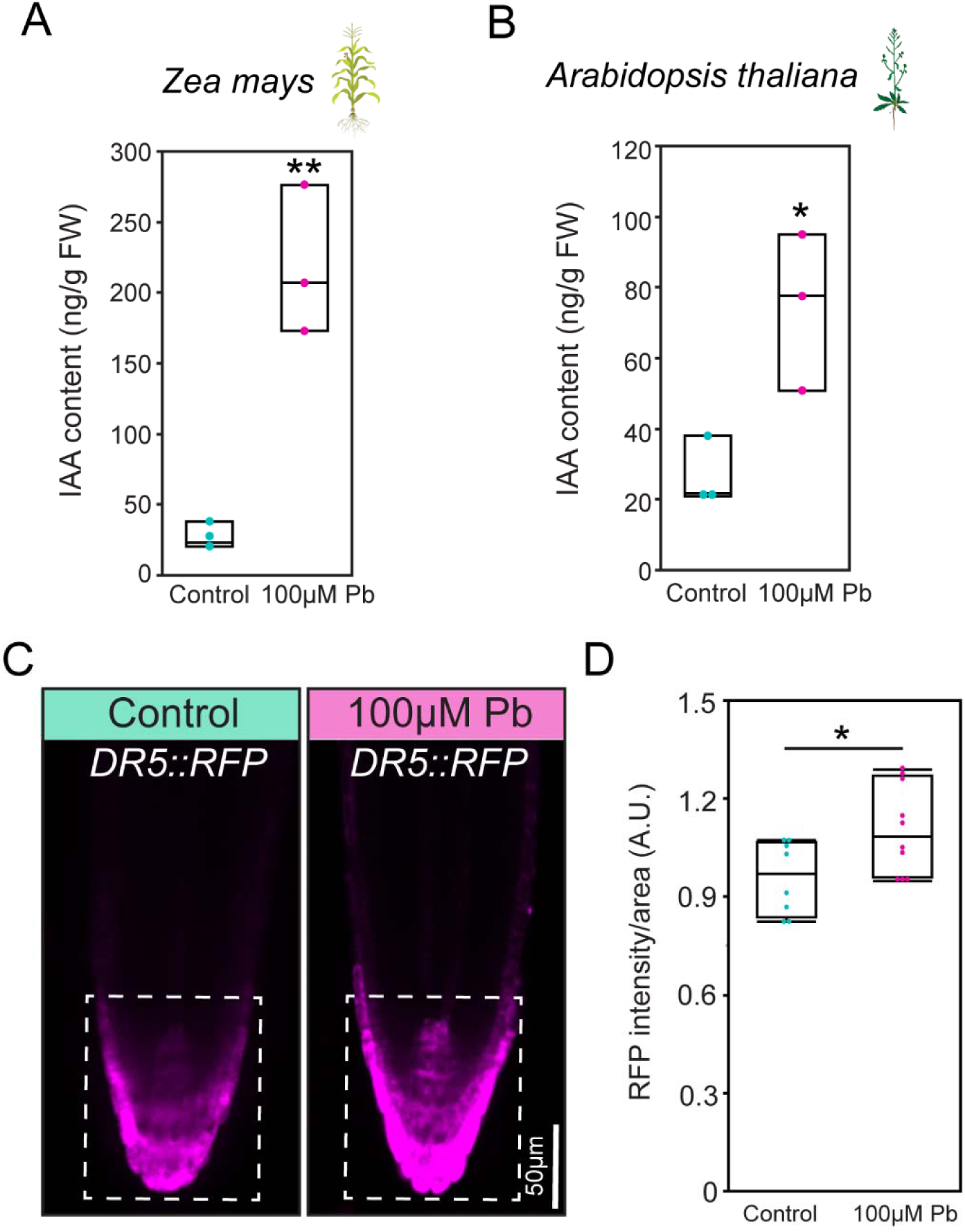
Auxin response is changed in response to lead. IAA quantification from maize (A) and Arabidopsis (B) root in absence and presence of 100µM Pb. (C) DR5::RFP signal in Arabidopsis root after 2 days of incubation in control and 100µM Pb plates. Scale bar = 50 µm. (D) Quantification of RFP signal from C. Box Plots show median values (center line), 25th to 75th interquartile range (box) and 1.5*interquartile range (whiskers). Student’s t-test was performed for A, B, and C. Asterisks represent the statistical significance between the means for each treatment. *P < 0.05, **P < 0.01, and ***P < 0.001.

**Table 2:** Hormone quantification from Arabidopsis root in absence (control) and presence of Pb. Higher than control is highlighted with cyan.

| Content<br>(ng/g FW) | *Arabidopsis thaliana* |  |
| --- | --- | --- |
|  | Control | 100µM Pb |
| **12OHJA** | 188.43 ± 70.22 | 254.68 ± 81.88 |
| **JA** | 575.57 ± 47.65 | 786.46 ± 35.24 (*) |
| **1-ACC** | 47.34 ± 2.53 | 60.62 ± 0.91 (*) |
| **ABA** | 184.10 ± 13.77 | 164.05 ± 5.98 |
| **GA53** | 44.55 ± 2.80 | 61.17 ± 5.59 |
| **GA12** | 65.50 ± 11.70 | 76.05 ± 18.03 |
| **IAA** | 45.19 ± 6.32 | 103.74 ± 23.97 (*) |
| **SA** | 80.34 ± 10.29 | 108.08 ± 11.59 |

Comparative transcriptomic analyses of root tips after 48h of Pb exposure revealed numerous auxin-related genes that were differentially expressed as compared to control treatment in both Arabidopsis and maize (Supplemental Figures S5 and S6). These included genes regulating auxin synthesis, transport, and downstream gene expression response. Collectively, these results implicate auxin as a critical regulator of the Pb-stress response.

### Lead alters auxin distribution in the primary root of Arabidopsis

While auxin was increased in both maize and Arabidopsis root tips, the localization and spatial distribution of auxin, rather than absolute levels are critical for downstream signaling and response. Therefore, the Arabidopsis DR5rev::erRFP (Marin et al. 2010) and proDR5::GUS (Ulmasov et al. 1997) auxin reporters were analyzed in the presence of Pb. This analysis localized the increased auxin to the columella cell regions of the meristem (Figures 3C and 3D, Supplemental Figure S7). Auxin-cytokinin crosstalk is a common regulator of root growth and development and was previously reported for Cd stress (Bruno et al. 2017). Our whole root phytohormone extraction protocol was not compatible with the detection of cytokinin, so we tested the response of cytokinin using the TCSn::GFP (Liu and Müller 2017) reporter in Arabidopsis. We found that Pb treatment substantially reduced the TCSn::GFP signal in columella cell regions consistent with a role for auxin-cytokinin crosstalk in Pb-responses (Supplemental Figure S8).

**Figure 3.**
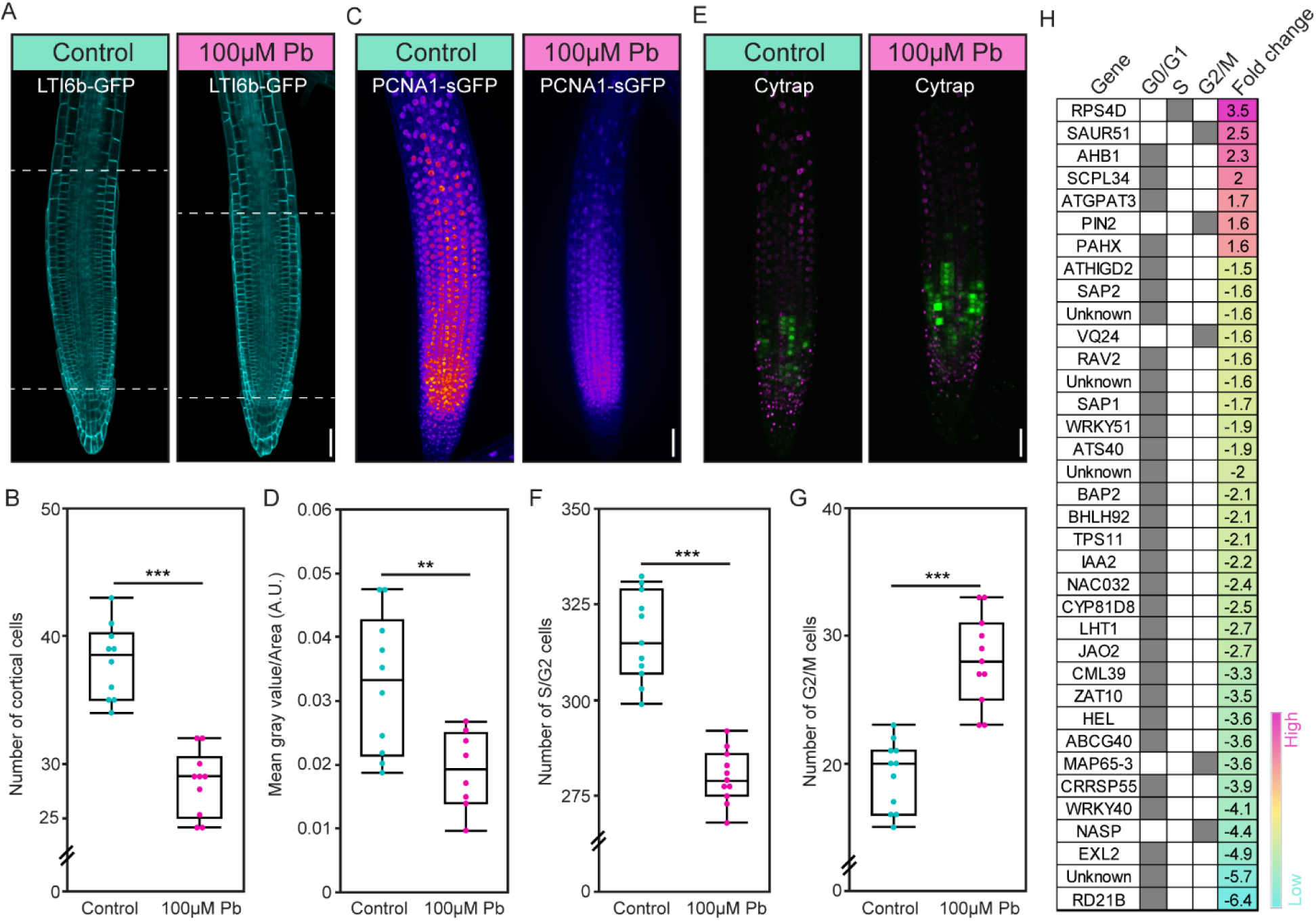
Lead alters cell cycle regulation in the primary root of Arabidopsis. LTI6b-GFP (A), PCNA1-sGFP (C), and Cytrap (E) signal in absence and presence of 100µM Pb. Scale bar = 50 µm. (B) Quantification of cortical cell numbers from A. (D) Quantification of gray value/area (A.U.) based on the PCNA1-sGFP signal in B. F and G highlights the quantification of the number of S/G2 and G2/M cells, respectively, in E. (H) Common differentially expressed genes in both Arabidopsis and maize along the cell-cycle specific expression pattern and fold change. Boxplots show median values (center line), 25th to 75th interquartile range (box) and 1.5*interquartile range (whiskers). Student’s t-test was performed for B, D, F, and G. Asterisks represent the statistical significance between the means for each treatment. *P < 0.05, **P < 0.01, and ***P < 0.001.

A brassinosteroid marker, proBZR1::BZR1-YFP (Tang et al. 2011), was used to confirm that Pb had no effect on brassinosteroid response as suggested by the hormone profiling (Supplemental Figure S9). There was no discernable difference in the proBZR1::BZR1-YFP localization or expression between the control and Pb treated plants. Together, these findings confirm Pb-induced altered auxin response in a columella-specific manner and highlight auxin-cytokinin crosstalk through hormone quantification and molecular reporter analysis.

### Lead inhibits primary root growth through cell cycle inhibition

Alteration of auxin in the root tip is reminiscent of a reduced root growth phenotype in various auxin mutants. We hypothesized that the Pb-induced auxin accumulation was responsible for root growth inhibition. Primary root growth is the outcome of both cell division and cell elongation (Rahman et al. 2007; Ashraf and Rahman 2019; Hazelwood et al. 2026). To determine which mechanism was inhibited under Pb, we first quantified the meristematic cell number in Arabidopsis roots grown in the presence and absence of Pb (Figure 2). The number of cortical cells, as marked by LOW TEMPERATURE INDUCED PROTEIN 6B-GREEN FLUORESCENT PROTEIN (LTI6b-GFP) expression, were significantly reduced in the Pb-treated root meristems (Figures 2A and 2B). In contrast, the epidermal cell length was not different between control and Pb-treated conditions (Supplemental Figure S10). These data indicate a reduction in cell cycle, and not cell elongation, is the cause of Pb-mediated root growth inhibition.

To further refine the cell cycle regulation, two cell cycle markers were analyzed. First, a cell cycle marker, PROLIFERATING CELL NUCLEAR ANTIGEN1-superfoldedGFP (PCNA1-sGFP) shows three distinct nuclear patterns corresponding with different phases of the cell-cycle– whole (G1/G2 phase), dotted (early S phase), and speckled (late S phase) (Yokoyama et al. 2016; Hazelwood and Ashraf 2024). Interestingly, these distinct nuclear patterns are not observed after the Pb treatment due to low expression of PCNA1-sGFP (Figures 2C and 2D), which indicates a reduction of earlier stages (G1/S/G2) of the cell cycle. A second cell cycle marker called Cell Cycle Tracking in Plants (Cytrap), which marks S/G2 and G2/M phases, was also used. Pb-treated roots showed a reduced number of cells at the S/G2 stage (Figures 2E-F), which corroborates with our observation from the PCNA1-sGFP experiment. And an increased number of cells at G2/M stage (Figures 2G) indicates a delay during mitosis with Pb-treatment.

Using the whole-root transcriptomic analyses, we took differentially expressed genes that were shared between both species and compared them against Arabidopsis cell cycle stage–specific single cell RNA sequencing datasets (Vukašinović et al. 2025). The analysis showed enrichment of differentially expressed genes specific to the G0/G1 stage when compared to the S or G2/M stages (Figures 2H). Root development markers (Marquès-Bueno et al., 2016), including the stele-specific *proSHR::H2B-2xmCherry*, endodermis-specific *proSCR::H2B-2xmCherry*, and quiescent center (QC)-specific *proWOX5::H2B-2xmCherry,* did not show any alteration in cell fate specification (Supplemental Figure S5) and suggests that cell identity during root development remains unchanged in the presence of Pb. Collectively, these results show that Pb inhibits the cell cycle progression which results in reduced root growth under Pb-stress.

### Pb causes altered gravitropic response in Arabidopsis and maize

Proper auxin distribution helps to maintain gravitropic responses at the root tip (Zhang et al. 2019b). As Pb alters auxin response in the root tip (Figure 3), we hypothesized that it may cause aberrant gravitropic response. To test the gravitropic response, we transferred 4-days-old Arabidopsis seedlings to control or 100μM Pb plates, incubated 12h to facilitate the Pb uptake, and rotated 90 degrees in anti-clockwise and clockwise directions (Figure 4). In both directions, we observed an aberrant gravitropic response at 3h, 6h, 9h, 12h, and 24h time points (Figures 4A-C). To test if this response is conserved across species, we followed up the Arabidopsis gravitropic response experiment with maize in the absence and presence of Pb and observed a similar slow gravitropic response of the maize root after Pb treatment (Figures 4D). Altogether, these experiments suggest that Pb alters a root-specific gravitropic response.

**Figure 4.**
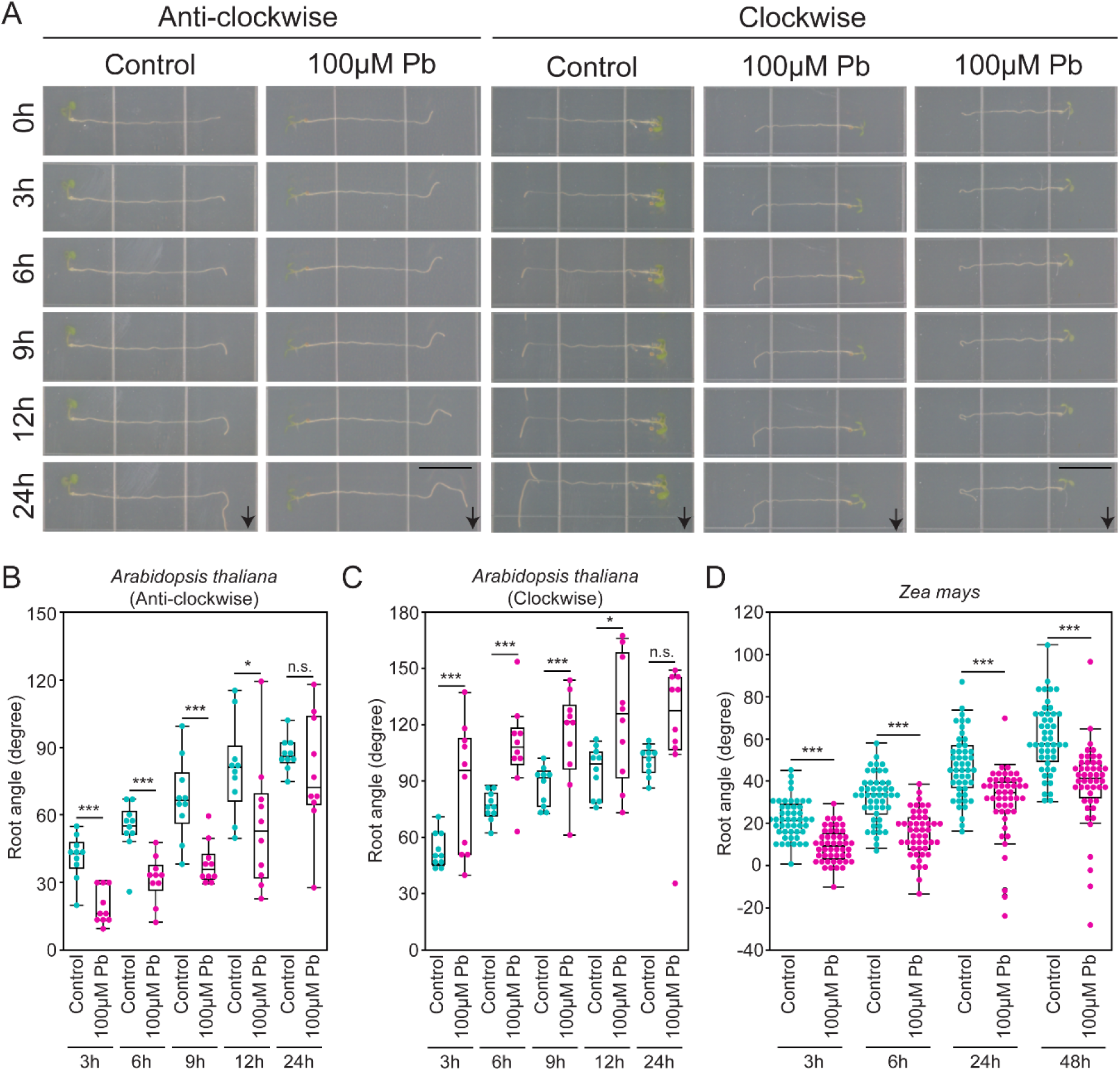
Lead alters gravitropic response in Arabidopsis and maize. (A) Anti-clockwise (left panel) and clockwise (right panel) gravitropic response in absence (control) and presence of 100µM Pb at 0, 3, 6, 9, 12, and 24h time points. Scale bare = 10mm. (B) Quantification of root angle after anti-clockwise gravistimulation in Arabidopsis. (C) Quantification of root angle after clockwise gravistimulation in Arabidopsis. (D) Quantification of root angle after clockwise gravistimulation in maize. Boxplots show median values (center line), 25th to 75th interquartile range (box) and 1.5*interquartile range (whiskers). Student’s t-test was performed for B, D, F, and G. Asterisks represent the statistical significance between the means for each treatment. *P < 0.05, **P < 0.01, and ***P < 0.001.

The fastest gravitropic response is manifested by the distribution of starch granules, also known as statolith, in the columella cells of the root tip (Zhang et al. 2019a). We aimed to image the starch granule distribution under control and Pb-treated conditions at earlier time points (0, 15, 30, and 60 minutes) after gravistimulation (Figure 5). To image the starch granules, we transferred 4-days-old Arabidopsis seedlings to the control and 100μM Pb plates, incubated 12h to facilitate the Pb uptake, rotated clockwise 90 degrees, and started to image at 0, 15, 30, and 60 minutes (Figure 5). We quantified each individual starch granule and determined their position as sedimented (bottom half of the cell towards the gravitropic orientation) and non-sedimented (upper half of the cell and opposite to the gravitropic orientation). We observed that less starch granules are sedimented at 0 minute time points indicating that the Pb treatment affects starch granule distribution even without gravistimulation. The fraction of non-sedimented starch granules significantly increased at 15 and 30 minute time points for Pb-treated roots as compared to the 0 minute time point (Figure 5). The fraction of non-sedimented starch granules remain the same between 30 and 60 minutes time points (Figure 5). It suggests that the major sedimentation issue is manifested within the first 30 minutes period. These data show that Pb causes alteration of starch granules distribution in the columella cells and consequently leads to aberrant gravitropic response.

**Figure 5.**
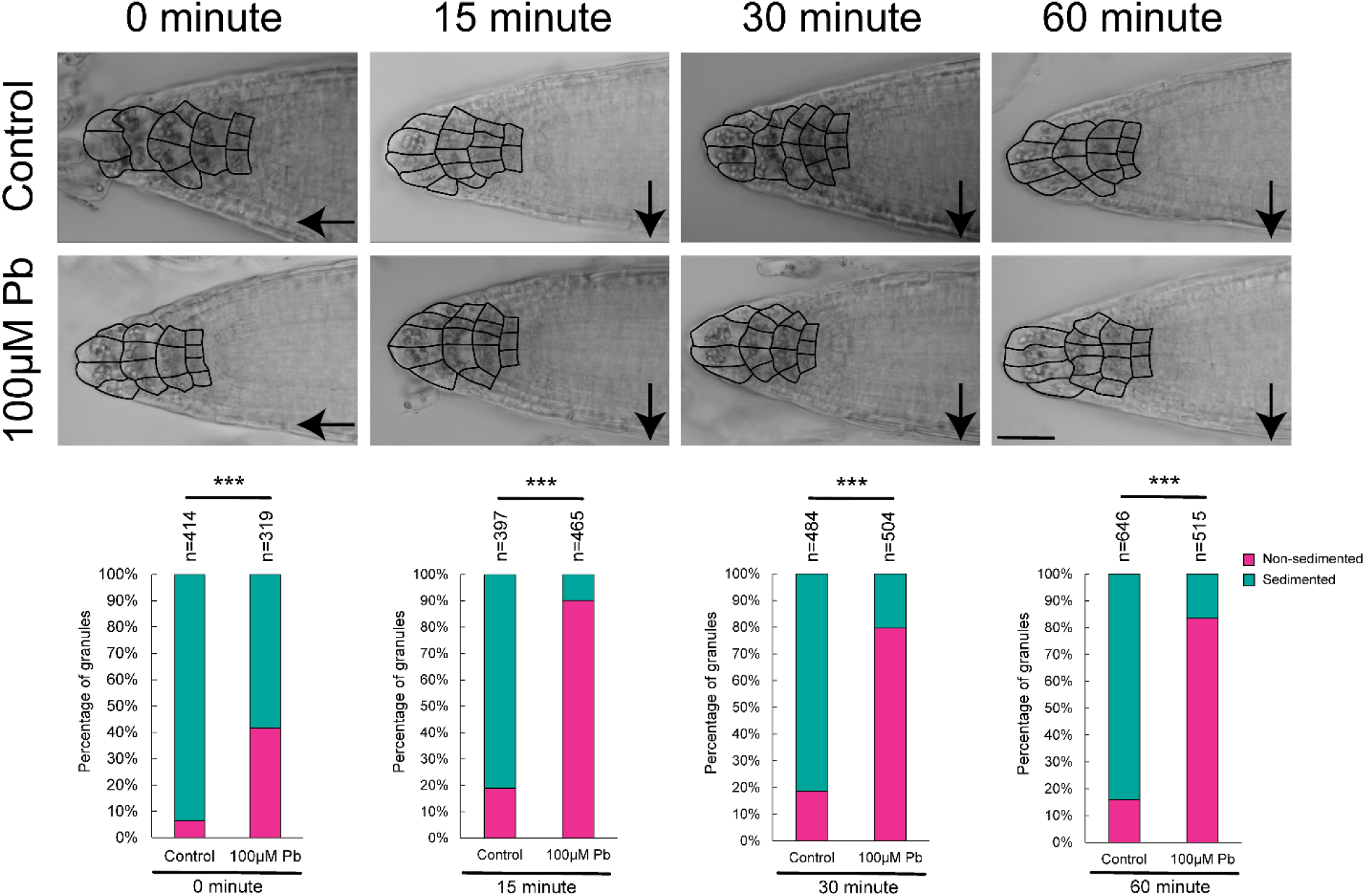
Lead causes aberrant starch granule distribution in columella cells. Representative images of columella cells after 0, 15, 30, and 60 minutes of gravistimulation in absence (control) and presence of 100µM Pb. At least 8-10 roots and 10-12 cells from root was considered to quantify the position of starch granules (N = number of starch granules). Starch granule sedimentation was compared in control and 100µM Pb conditions using a Fisher’s exact test. *P < 0.05, **P < 0.01, and ***P < 0.001.

### Pb-induced altered auxin distribution in the root tip leads to aberrant gravitropic response

The impaired statolith sedimentation should disrupt the spatial distribution of auxin upon gravistimulation. To test this, we analyzed the auxin ratiometric reporter line, R2D2 (Liao et al. 2015) during the pre- and post-gravistimulation in the absence and presence of Pb (Figure 6). For this purpose, we transferred 4-days-old R2D2 seedlings to the control and 100μM Pb plates, incubated 12h to facilitate the Pb uptake, and imaged the R2D2 reporters simultaneously for DII-n3xVenus and mDII-ntdTomato at the 0h time point (Figure 6A). We quantified the Venus and tdTomato signal along with the area (meristematic region with similar Z stacks across images). Then, we calculated the ratio of DII/mDII for control and 100μM Pb. We found a lower DII/mDII ratio value in Pb-treated roots, which indicates a higher auxin response (Figure 6A) and corroborates our previous observations (Figure 3).

**Figure 6.**
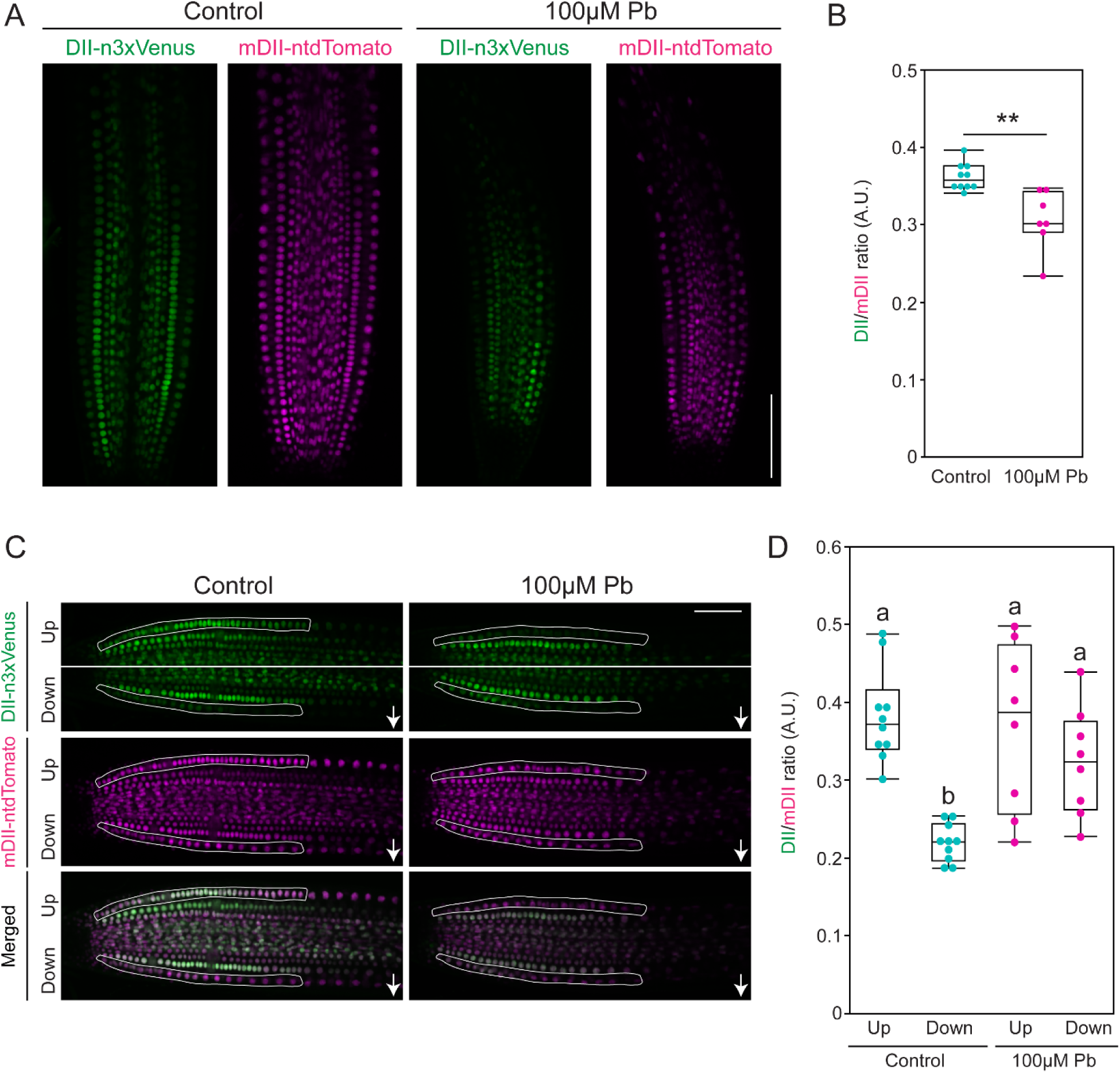
Lead alters spatial distribution of auxin in the primary root. (A) Visualization of DII-n3xVenus (green) and mDII-ntdTomato (magenta) using R2D2 in absence (control) and presence of 100µM Pb. 4-days-old R2D2 seedlings were incubated in absence (control) and presence of 100µM Pb for 12h and imaged. Scale bar = 100µm. (B) Quantification of DII/mDII from (A). Boxplots show median values (center line), 25th to 75th interquartile range (box) and 1.5*interquartile range (whiskers). Student’s t-test was performed. Asterisks represent the statistical significance between the means for each treatment. *P < 0.05, **P < 0.01, and ***P < 0.001. (C) Visualization of DII-n3xVenus (green) and mDII-ntdTomato (magenta) using R2D2 after 30 minutes of gravistimulation in absence (control) and presence of 100µM Pb. The white arrow indicates the direction of gravistimulation. The region of interest is drawn with white markers for both upper and lower sides of the root. Scale bar = 50µm. (D) Quantification of DII/mDII for both upper (Up) and lower (Down) side of the root after gravistimulation from (C). Statistical test is performed based on Tukey’s Honest test. Groups labeled with the same letter are not statistically different from each other (alpha = 0.05). Boxplots show median values (center line), 25th to 75th interquartile range (box) and 1.5*interquartile range (whiskers).

We followed up the spatial auxin distribution imaging of R2D2 along with gravistimulation. A positive gravistimulation requires asymmetric auxin distribution between upper and lower parts of the root, where auxin is more accumulated on the lower side (Brunoud et al. 2012; Liao et al. 2015). We rotated the R2D2 seedlings 90 degrees for 30 minutes and imaged simultaneously for DII-n3xVenus and mDII-ntdTomato (Figure 6C). We quantified the Venus and tdTomato signal in the epidermal cell layers for the upper and lower part of the root and calculated the DII/mDII ratio for both upper and lower sides. We observed a reduction in DII/mDII ratio for the lower side of the root after the gravistimulation in the control treatment indicating a positive gravitropic response (Figure 6D). Interestingly, a sporadic DII/mDII ratio was observed after 30 minutes of gravistimulation of Pb-treated roots (Figure 6D). These results indicate that spatial auxin distribution is altered due to Pb toxicity and consequently, the root fails to maintain a positive gravitropic response.

### Pb inhibits PIN3 trafficking in the columella cells

Next, we searched for the molecular and cellular target for Pb to alter the auxin response. We took a reverse genetics approach to answer this question. We tested a series of auxin mutants, including auxin biosynthesis (*wei2-2*, *wei8-2*, and *wei2-1 wei7-1*), signaling (*tir1-10* and *axr1-3*), and transport (*pin1-3*, *eir1-1*, *pin3-4*, *pin4-3*, *pin7-3*, and *aux1-7*) mutants (Supplemental Figure S12). If auxin is critical for the Pb-mediated response, we would expect the mutants to be insensitive to Pb and maintain root growth in the presence of Pb. The biosynthesis and signaling mutants we chose disrupted all auxin production and signaling, and all were insensitive to Pb, which is consistent with a role of auxin in mediating the response. Although, the double *wei2-1 wei7-1* mutant is already significantly shorter making a response difficult to distinguish.

In contrast, we screened several different transporters, and identified the *pin3-4* mutant as the only transporter that was resistant to Pb treatment for primary root growth inhibition (Supplemental Figure S12). The *pin3-4* mutant complemented line responded normally to Pb-induced root growth inhibition, demonstrating the specificity of this response (Figure 7).

**Figure 7.**
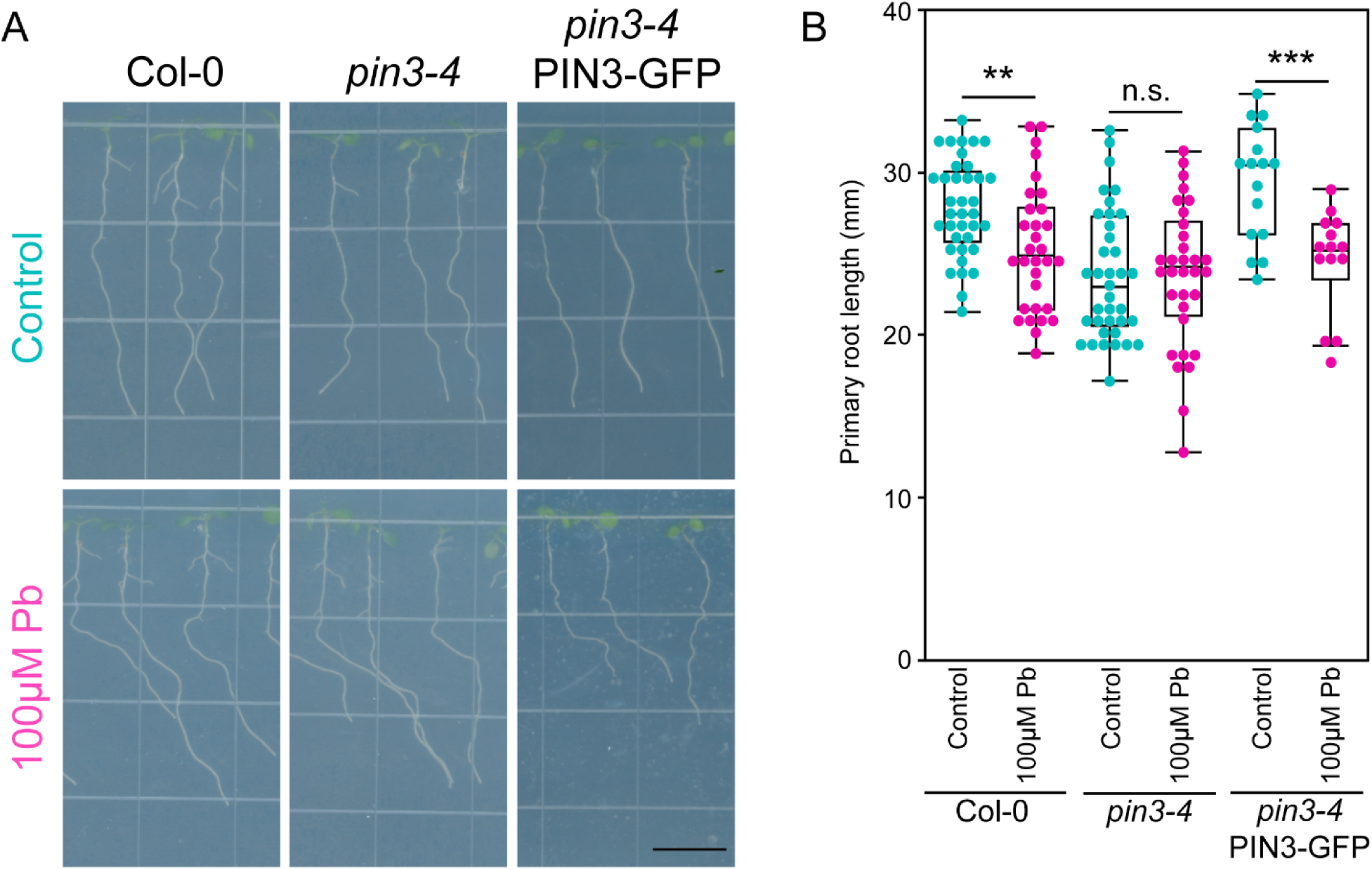
PIN3 is the genetic regulator of lead response in Arabidopsis root. (A) 3-days-old Col-0, *pin3-4*, and *pin3-4* PIN3-GFP lines were transferred to control and 100µM Pb containing plates to observe the primary root growth for 2 days. Scale bar = 10mm. (B) Quantification of primary root growth from (A). Boxplots show median values (center line), 25th to 75th interquartile range (box) and 1.5*interquartile range (whiskers). Student’s t-test was performed. Asterisks represent the statistical significance between the means for each treatment. *P < 0.05, **P < 0.01, and ***P < 0.001.

The PIN3 auxin transporter localizes at the columella cell regions of the root tip (Friml et al. 2002). We tested the PIN3 localization, using *proPIN3::PIN3-GFP*, in the absence and presence of 100μM Pb. Interestingly, the overall PIN3 localization pattern and intensity (mean gray value/area) remained the same after Pb treatment (Figures 8A and 8B), which is consistent with the transcriptomic results which did not identify PIN3 as differentially expressed. However, auxin transporter proteins require intracellular trafficking, which can be disrupted by application of BFA (Geldner et al. 2003). We incubated control and Pb-treated PIN3-GFP seedlings with 40μM BFA for 60 minutes and imaged the columella cells (Figure 8C). We quantified the number of PIN3-positive BFA bodies formed per columella cell and found a reduced number of BFA bodies appeared after the Pb treatment (Figure 8D). This data suggests that Pb inhibits PIN3 protein trafficking in the columella cells, which disrupts auxin flow to mediate local Pb-responses and inhibit root growth and gravitropic responses.

**Figure 8.**
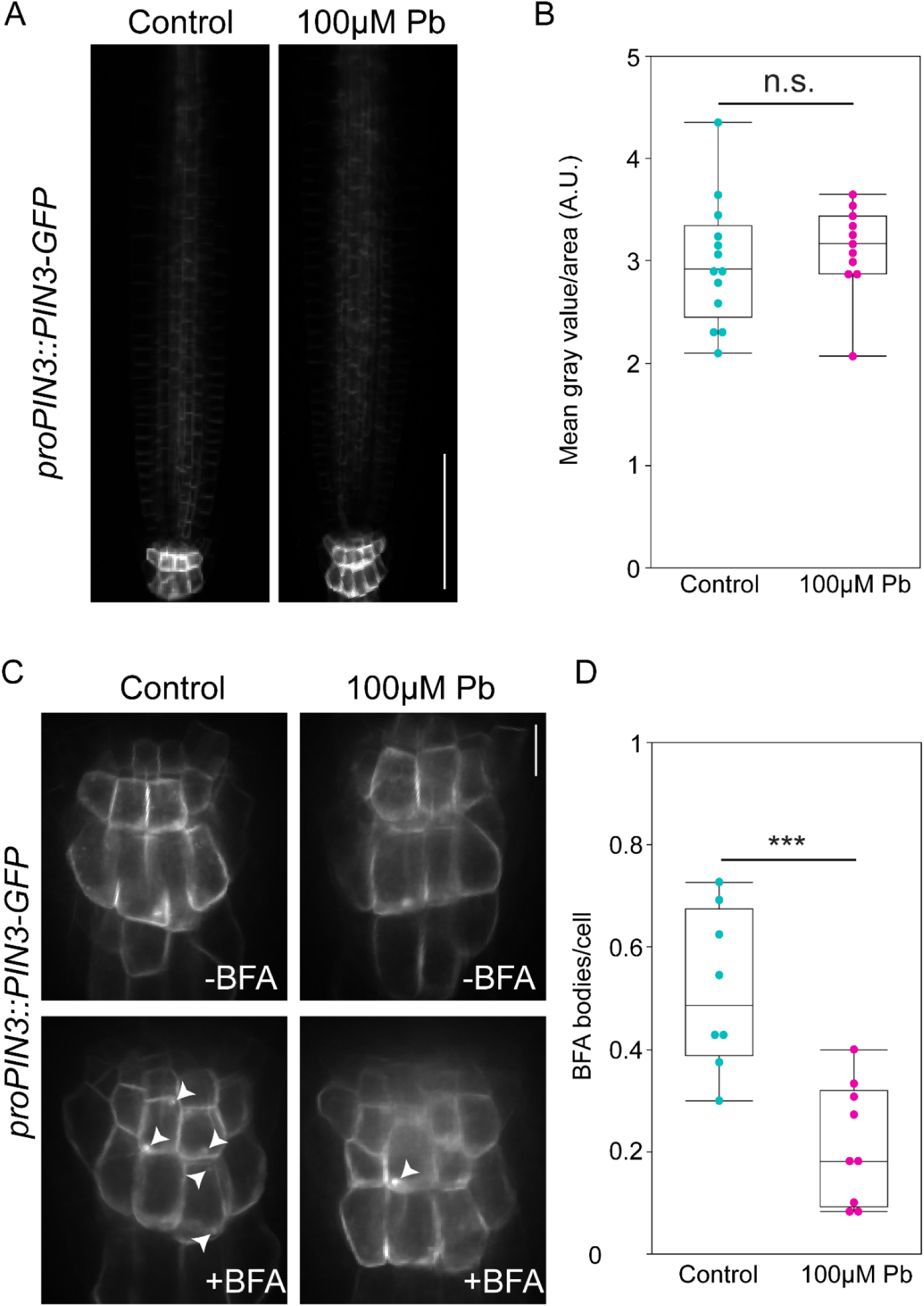
Lead inhibits PIN3 protein trafficking in the columella cells. (A) 3-days-old *pin3-4 proPIN3::PIN3-GFP* line was transferred to control and 100µM Pb containing plates and incubated for 2 days. Scale bar = 50µm. (B) Quantification of GFP intensity/area from (A). Boxplots show median values (center line), 25th to 75th interquartile range (box) and 1.5*interquartile range (whiskers). Student’s t-test was performed. Asterisks represent the statistical significance between the means for each treatment. *P < 0.05, **P < 0.01, and ***P < 0.001. (C) Visualization of columella cells of *pin3-4 proPIN3::PIN3-GFP* line using 60x (oil) objective after incubating two days in control and 100µM Pb conditions. 40μM BFA was used for 60 minutes treatment before imaging. White arrow head indicates visible BFA bodies. Scale bar = 50µm. (D) Quantification of BFA bodies per cell from (C). Boxplots show median values (center line), 25th to 75th interquartile range (box) and 1.5*interquartile range (whiskers). Student’s t-test was performed. Asterisks represent the statistical significance between the means for each treatment. *P < 0.05, **P < 0.01, and ***P < 0.001.

## Discussion

In this study, we show that two evolutionarily distant plant species, Arabidopsis and maize (Sharma et al. 2022), have conserved responses to toxic metal contamination, but that the responses to different toxic metals are distinct. Arabidopsis roots exhibited distinct altered root morphology when treated with a different toxic metal. The hormone profile responses of maize were also distinct between the different toxic metals. These results demonstrate the importance of studying plant responses to individual toxic metals and developing targeted bioremediation strategies. A plant ideotype for bioremediation can explore the soil to remove toxic metals, while locally compartmentalizing the metals for removal from the soil.

For Pb, which is arguably the most widespread toxic metal contaminant (Jaishankar et al. 2014), we have shown that it inhibits root growth and alters gravitropic response by inhibiting the protein trafficking of auxin transporter PIN3 in columella cells resulting in increased auxin accumulation in the root. Both of these responses are critical for soil exploration, which is key to effective bioremediation (Hou et al. 2020). If roots are unable to grow and explore the soil, they will be highly ineffective at remediation. Thus, plants that can maintain root growth in the presence of lead are critical for future bioremediation efforts. Targeting PIN3 and its response to Pb will be an effective breeding strategy to improve growth responses under toxic Pb environments.

Although a set of previous studies demonstrated the Pb-induced root growth inhibition in Arabidopsis (Ortiz-Luevano et al. 2021; Zheng et al. 2021) and maize (Jiang and Liu 2000) roots, the mechanistic explanation was unknown. In this study, we took a comprehensive cell biology approach by utilizing Cytrap and PCNA1-GFP cell cycle markers. Quantitative measurement of cell cycle phases indicated that Pb toxicity causes inhibition at G1 phase. Previous studies only used CYCB1;1-GUS after the Pb incubation (Zheng et al. 2021) and as a result the cell cycle phase specific effect was not detected. Our study also analyzed the differentially expressed cell cycle genes resulting from Pb toxicity and analyzed the cell cycle phase specificity of these target genes from the single cell RNA sequencing data (Nolan et al. 2023). The majority of these differentially expressed genes in Arabidopsis and maize are G1-phase-specific. Both the quantitative cell biology and comparative transcriptomics approaches strongly suggested a cell cycle phase specific inhibition by Pb toxicity. The effects on cell cycle regulators are likely due to altered auxin distribution in the root tip.

In a previous study, a high concentration of Pb exposure was demonstrated to affect the DR5::GUS signal in Arabidopsis (Wang et al. 2015), but the mechanistic explanation of auxin response was not demonstrated. In this study, we used a comprehensive approach to measure major phytohormones and their derivatives from both Arabidopsis and maize roots. This quantitative approach not only shows the increased auxin content in both Arabidopsis and maize but also provides the largest phytohormone dataset in the presence of toxic metals. Previous studies have measured auxin levels in the maize shoot after Pb treatment and showed that it accumulates similar to what we showed occurs in the root. Interestingly, the Pb treatment and subsequent auxin accumulation in the shoot results in increased growth of the shoot (Małkowski et al. 2020). This indicates that auxin accumulates throughout the plant after Pb treatment but has opposite effects on growth in the root and shoot of the plant. It would be interesting to test if Pb affects gravitropic responses in the shoot similar to what was observed in the root.

Using quantitative live cell imaging, we have demonstrated that altered auxin response causes altered asymmetric auxin distribution and hampers the gravitropic response under Pb-stress (Figure 9). Finally, we identified columella-specific PIN3 as a molecular and cellular target during Pb toxicity and opens the door for future studies for gene editing and improved bioremediation in a wide range of crop plants (Figure 9).

**Figure 9.**
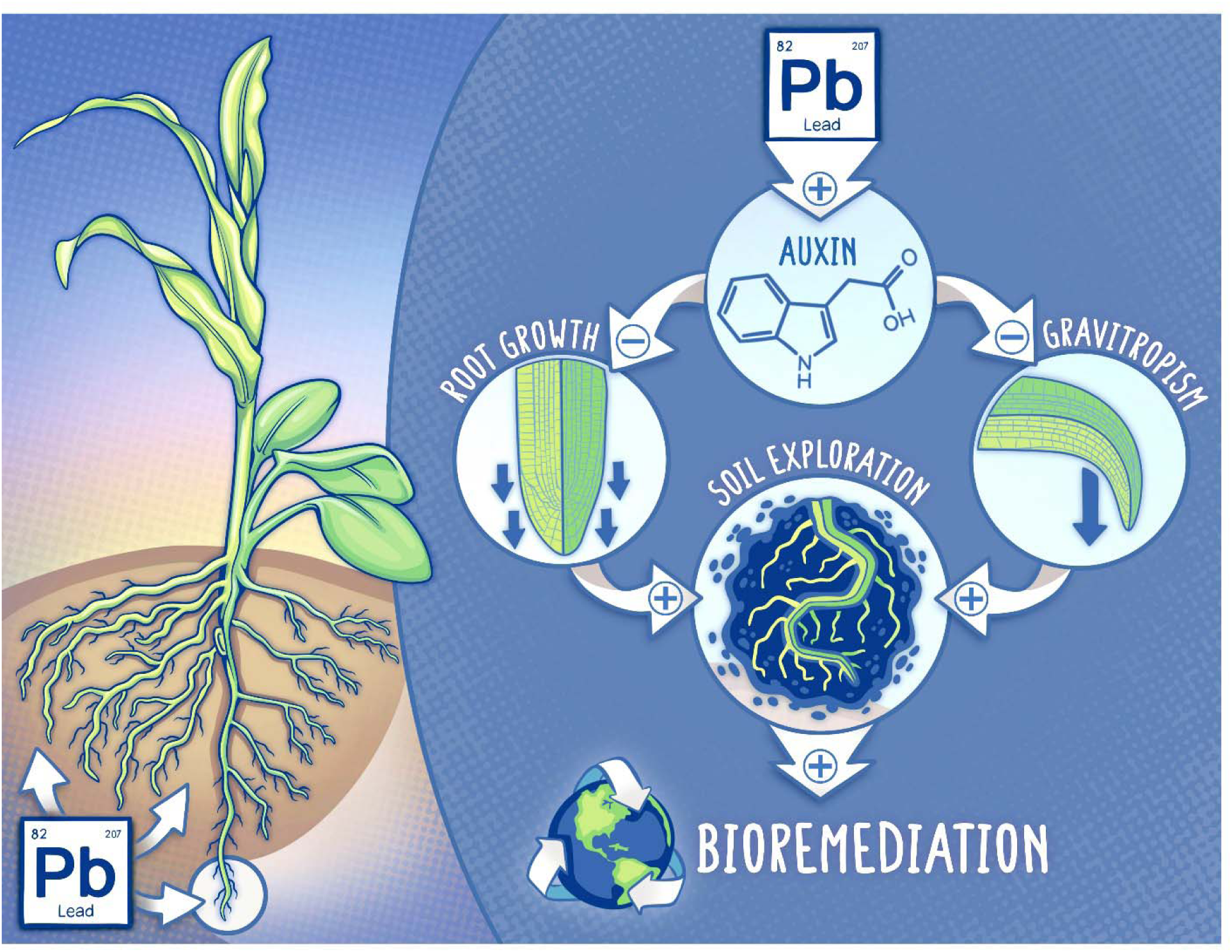
Graphical abstract of the study. Pb alters auxin in both Arabidopsis and maize root, and consequently affects the primary root growth and gravitropic response negatively. Both primary root growth and proper gravitropic responses are required for soil exploration.

## Materials and Methods

### Plant materials

Columbia-0 (Col-0) and B73 were used as the wild types for Arabidopsis and maize, respectively. Transgenic reporter lines used in this study were reported previously: LTI6b-EGFP (Kurup et al. 2005), PCNA1-sGFP (Yokoyama et al. 2016), Cytrap (Yin et al. 2014), *DR5rev::erRFP* (Marin et al. 2010), *DR5::GUS* (Ulmasov et al. 1997), R2D2 (Liao et al. 2015), *TCSn::GFP* (Liu and Müller 2017), *proBZR1::BZR1-YFP* (Tang et al. 2011), *proSHR::H2B-2x mCherry* (Marquès-Bueno et al. 2016), *proSCR::H2B-2xmCherry* (Marquès-Bueno et al. 2016), and *proWOX5::H2B-2x mCherry* (Marquès-Bueno et al. 2016). Auxin mutants and reporter, including *pin1-3* (CS69065), *eir1-1* (CS8058), *pin3-4* (CS9363), *pin4-3* (CS9368), *pin7-3* (CS9367), *aux1-7* (CS3074), *axr1-3* (CS3075), *axr1-12* (CS3076), *wei2-2* (CS16398), *wei2-1 wei7-1* (CS16399), *tir1-10* (CS69649), *wei8-2* (CS16408), pPIN3:PIN3-GFP (CS69618), R2D2 (CS2105637) were obtained from ABRC (https://abrc.osu.edu/).

### Plant growth conditions

Col-0 seeds were surface sterilized with 1mL 70% ethanol for 10 minutes, washed twice with 1mL autoclaved ultrapure water inside the laminar flow hood, and placed on the surface of ½ Murashige and Skoog (MS) media (BioWorld; Cat.# 30630058) containing 1% sucrose (BioWorld; Irving, TX) and 1% agar (BioWorld) in square petri dishes (Simport; Bernard-Pilon Beloeil, Canada). Plate openings were sealed with ½ inch thick micropore tape (Amazon; Seattle, WA), covered with aluminum foil, and kept at 4°C for 2 days. After 2 days, plates were placed vertically under constant light conditions at 22°C (Hazelwood et al., 2025).

B73 maize seeds were treated with 10% bleach for 10 mins and rinsed five times. Seeds were then transferred to petri dishes with germination paper with water and placed in an incubator in the dark at 29 °C for two days. After 48 hrs., germinated seedlings were transferred to plates with 2% agar (Becton Dickinson, Franklin Lakes, NJ; Cat. no. DF0812-07-1) and with or without toxic metal treatment. Plates were imaged at the specified time point using a flatbed scanner.

### Chemical treatment

Lead (II) nitrate (Sigma-Aldrich St. Louis, MO; Cat. no. 10099-74-8), Sodium (meta)arsinate (Sigma-Aldrich St. Louis, MO; Cat. no. 7784-46-5), Potassium chromate (Sigma-Aldrich St. Louis, MO; Cat. no. 7789-00-6), and Cadmium carbonate (Sigma-Aldrich St. Louis, MO; Cat. no. 10108-64-2) were dissolved in water to prepare the stock solution. For treating seedlings, mentioned final concentration was obtained by mixing with either ½ Murashige and Skoog (MS) media. For the protein trafficking inhibition experiment, Brefeldin A (BFA) (Sigma-Aldrich St. Louis, MO; Cat. no. B5936) was dissolved in DMSO and performed the experiment in the mentioned concentration.

### Gravitropic assay

4-days-old Arabidopsis seedlings were transferred to control and 100μM Pb containing plates, incubated for 12h, and rotated for the gravitropic response and start granules imaging. B73 maize seeds were treated with 10% bleach for 10 mins and rinsed five times. Seeds were then transferred to petri dishes with germination paper with water and placed in an incubator in the dark at 29 °C for two days. After 48 hrs., germinated seedlings were transferred to plates with 2% agar (Becton Dickinson, Franklin Lakes, NJ; Cat. no. DF0812-07-1) and with or without 100µM lead (II) nitrate (Sigma-Aldrich St. Louis, MO; Cat. no. 10099-74-8). Seedlings were then allowed to acclimate to the new plate for 48hrs when after they were rotated 90 degrees in the clockwise direction. Images of the plates were taken 0hr, 3hrs, 6hrs, 24hrs, and 48 hrs after rotating using a flatbed scanner. Images were analyzed using the SmartRoot software in ImageJ (Lobet et al. 2011).

### Phytohormone quantification

For LC MS/MS quantification of endogenous phytohormones about 100 mg of Arabidopsis or maize root tissue was ground in liquid nitrogen and an extraction solvent of acetonitrile was added along with a 1 µL spike-in each of deuterated d5-indole-3-acetic acid, d4-salicylic acid, d5-jasmonic acid, d6-abscisic acid, d2-gibberellic acid^53^, and d4-1-aminocyclopropane-1-carboxylate (1 µg/µL) internal standard. Samples were vortexed for 10 mins at 3,000 RPM and then incubated for 1 hr at -20°C. The samples were then centrifuged for 20 mins at 15,900 relative centrifugal force (RCF). Supernatant was transferred to a new tube and speed-vacuumed overnight at 45°C. The dried pellet was resuspended in 1.5 mL of 80% acetonitrile and vortexed for 3 mins at 3,000 RPM. Liquid was transferred to a new tube containing 10 mg graphitized carbon black and vortexed for 3 mins at 3,000 RPM. The samples were then centrifuged for 2 mins at 15,900 RCF. The supernatant was transferred to a new tube and filtered through a 0.2 µm syringe filter and speed-vacuumed for 6 hrs at 45 °C. Dried samples were stored at -20 °C until ready to be loaded on the LC MS/MS. Dried samples were then resuspended in 70 µL of 5% acetonitrile by vortexing at 3,000 RPM for 30 mins. A total of 20 µL was injected into an Agilent (Santa Clara, CA) 1260 Infinity II LC system coupled with an Agilent Ultivo MS/MS triple quadrupole. The LC and MS/MS method for measuring polar and nonpolar hormone metabolites was previously described (Dilkes and Best 2025a). The method for measurement of sterol and brassinosteroid intermediates was previously described (Dilkes and Best 2025b).

### GUS staining

GUS staining was performed based on a previously described method (Ashraf and Rahman 2019). Seedlings were transferred to GUS staining buffer (100 mM sodium phosphate, pH 7.0, 10 mM EDTA, 0.5 mM potassium ferricyanide, 0.5 mM potassium ferrocyanide and 0.1% Triton X-100) containing 1mM X-gluc and incubated at 37°C in the dark for 3h. The roots were imaged with a Leica (Wetzlar, Germany) DM 2000 LED compound microscope.

### Microscopy

Epidermal cell length was observed using a 20x objective lens attached with a Leica DM 2000 LED compound microscope setup and the images were taken by Leica MC170 HD camera attached with the Leica DM 2000 LED. *DR5::GUS* images were taken using a similar microscope setup. Starch granules imaging in Arabidopsis was performed using a 40x objective lens attached with AmScope T490 Series Simul-Focal Biological Trinocular Compound Microscope and images were captured using a 18MP USB 3.0 C-mount Camera and AmScope software.

Fluorescent images of live Arabidopsis root meristem of marker lines (*DR5rev::erRFP*, DII-Venus, *TCSn::GFP*, *proBZR1::BZR1-YFP*, *proSHR::H2B-2x mCherry*, *proSCR::H2B-2xmCherry*, and *proWOX5::H2B-2x mCherry*) were imaged using a Nikon (Minato City, Japan) Ti-E-PFS inverted spinning-disk confocal microscope equipped with a 20× objective. The system is outfitted with a Yokogawa (Musashino, Japan) CSU-X1 spinning disk unit, a self-contained 4-line laser module (excitation at 405, 488, 561, and 640 nm), and Andor (Belfast, Northern Ireland) iXon 897 EMCCD camera. 488nm and 561nm excitation were used for GFP/YFP/Venus and RFP/mCherry/PI, respectively. Fluorescent images were acquired using the Nikon NIS-Elements software and processed using ImageJ software (<u>imagej.nih.gov/ij/</u>) (Schindelin et al. 2012).

LTI6b-GFP, PCNA1-sGFP, Cytrap, and PIN3-GFP images were acquired using an Olympus (Evident; Shinjuku, Japan) IXplore SpinSR system, equipped with an inverted IX83 microscope (Evident), a CSUW1-Sora spinning disk confocal unit (Yokogawa) and a Hamamatsu (Shizuoka, Japan) ORCA Fusion BT camera. Fluorescence images were obtained with a 20x or 63x (oil) objective (UPLXAPO 20X/0.8NA). Fluorescence confocal images were acquired using software cellSens (Evident).

### Microscopic image quantification

For the intensity quantification of R2D2, the epidermis cell layer from the meristematic regions of the upper and lower part of the root after gravistimulation were considered. From each section, both DII-n3xVenus and mDII-ntdTomato signals were quantified. The intensity ratio of DII-n3xVenus/mDII-ntdTomato was calculated and plotted.

### RNA-sequencing

Three replicates of maize roots treated with and without 100µM Pb for 2 days were collected, flash frozen with liquid nitrogen, and total RNA was extracted using E.Z.N.A. Plant RNA Kit from Omega Bio-Tek (Norcross, GA). Poly-A enriched RNA-seq libraries were prepared by using Vazyme VAHTS Universal V10 RNA-seq Library Prep Kit and sequenced at 150-bp PE on an Element AVITI instrument by AmpSeq (Gaithersburg, MD) (https://www.ampseq.com/). Sequenced reads were trimmed and filtered by Trimmomatic (v0.39) (Bolger et al. 2014). Filtered reads were then aligned to the B73 reference genome (v5.0.52) by hisat2 (v2.2.1) (Kim et al. 2019; Woodhouse et al. 2021). Aligned reads were then calculated into counts tables by htseq (v2.0.2) (Anders et al. 2015). Differential expression was calculated using DESeq2 (v1.47.3) (Love et al. 2014). Four replicates of Arabidopsis plants were treated with and without 100µM Pb for 2 days and then were collected and analyzed as described previously, however these reads were aligned to the Arabidopsis reference genome (vTAIR10.60) (Lamesch et al. 2012) (Lamesch et al., 2012). Sequencing information and statistics are described in Supplemental Table 1.

### Statistical analysis

The raw data for each quantification was imported to statistical software JMP (Cary, NC) Pro18 for generating graphs and performing statistical tests.

## Data and materials availability

RNA-seq data are available on NCBI SRA under BioProject PRJNA1255621.

## Funding

The research at Ashraf lab is funded by the NSERC Discovery grant (RGPIN-2025-04277) Canada Foundation for Innovation (CFI) John R. Evans Leaders Fund (JELF), BC Knowledge Development Fund (BCKDF), and start-up grant provided by the University of British Columbia. Olivia S. Hazelwood is supported by the Hugo E Meilicke Memorial Fellowship, and Frances Chave Memorial Graduate Scholarship. Grace Delpit and Scottie Mello were supported by the ASPB SURF program in the summer of 2024 and 2026, respectively. Scottie Mello was supported by the Work Learn International Undergraduate Research award by the University of British Columbia.

This work was funded by the United States Department of Agriculture (USDA)-Agricultural Research Service Project number 5070-21220-046-000-D to NBB. This research used resources provided by the SCINet project and/or the AI Center of Excellence of the USDA Agricultural Research Service, ARS project numbers 0201-88888-003-000D and 0201-88888-002-000D. The use of trade name, commercial product or corporation in this publication is for the information and convenience of the reader and does not imply an official recommendation, endorsement, or approval by the USDA or the Agricultural Research Service for any product or service to the exclusion of others that may be suitable. USDA is an equal opportunity provider and employer.

## Acknowledgements

The authors thank Masaaki Umeda (Nara Institute of Science and Technology), Sachihiro Matsunaga (Tokyo University of Science), Abidur Rahman (Iwate University), and Zhiyong Wang (Carnegie Institution for Science, Stanford) for sharing Cytrap (Yin et al. 2014), *pAtPCNA1::AtPCNA1-sGFP* (Yokoyama et al. 2016), LTI6b-EGFP (Kurup et al. 2005), and *proBZR1::BZR1-YFP* (Tang et al. 2011), respectively, seeds. Authors thank Miki Fujita and EunKyoung Lee of UBC Bioimaging Facility (RRID: SCR_021304) for their kind support.

## Authors contributions

M.A.A., N.B.B., and E.E.S. adopted the project and idea and supervised. J.Z., S.M., O.S.H., G.D., N.B.B., and M.A.A. performed the experiments and analyzed the data. M.A.A., N.B.B., and E.E.S. wrote the manuscript. All authors agreed to the final version of the manuscript.

## Conflicts of interests

E.E.S. is co-founder of Izbe Innovations LLC. The company was not involved in the design, conduct, analysis, or funding of this work, and the subject matter of this publication is unrelated to the company’s business activities.

---

**Supplemental figure 1.**
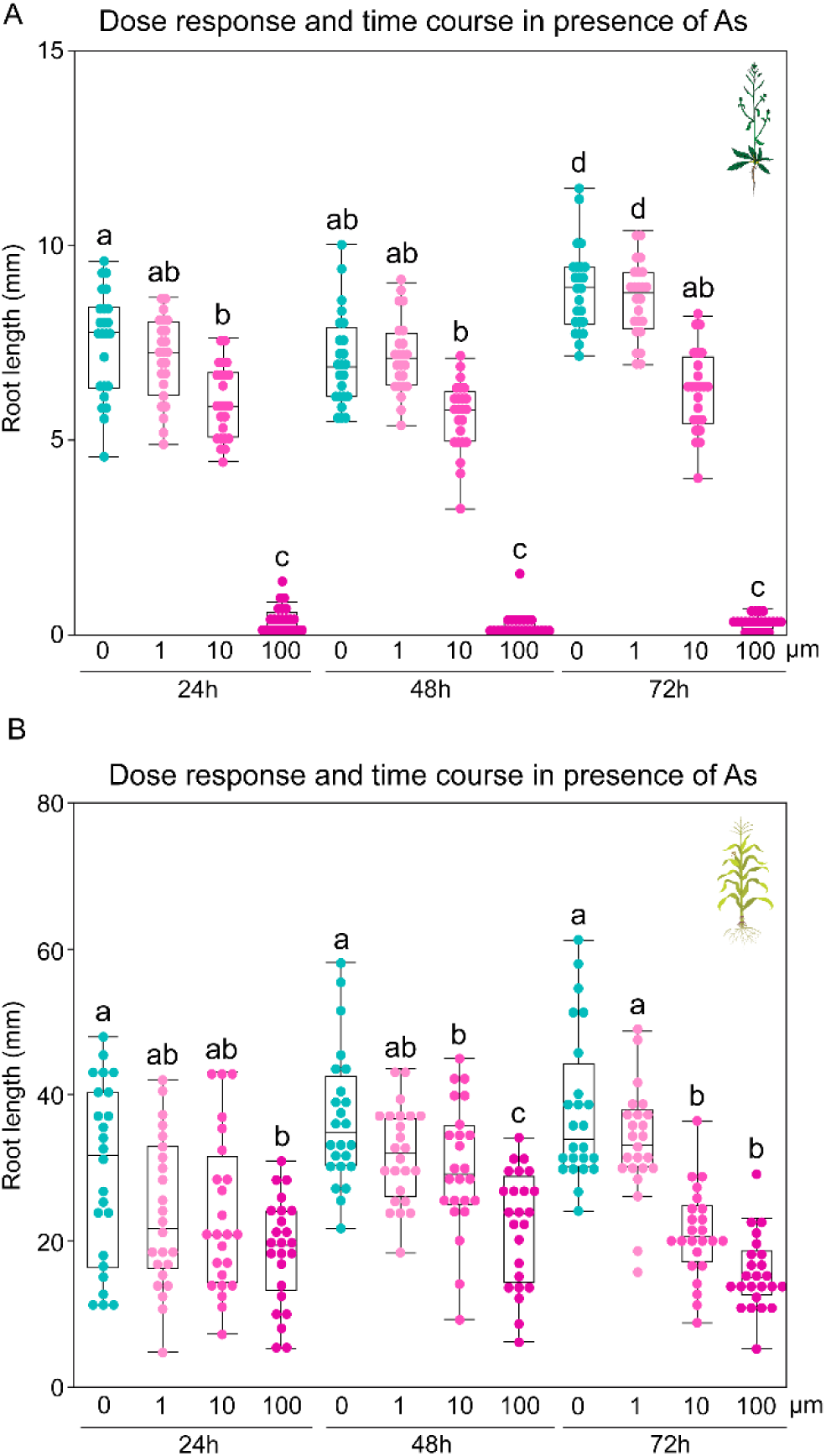
Time course and dose response of As in Arabidopsis and maize. (A) Quantification of primary root growth of Arabidopsis after incubating at control, 1µM As, 10µM As, and 100µM As conditions for 24h, 48h, and 72h time points. (B) Quantification of primary root growth of maize after incubating at control, 1µM As, 10µM As, and 100µM As conditions for 24h, 48h, and 72h time points. Statistical test is performed based on Tukey’s Honest test. Groups labeled with the same letter are not statistically different from each other (alpha = 0.05). Boxplots show median values (center line), 25th to 75th interquartile range (box) and 1.5*interquartile range (whiskers).

**Supplemental figure 2.**
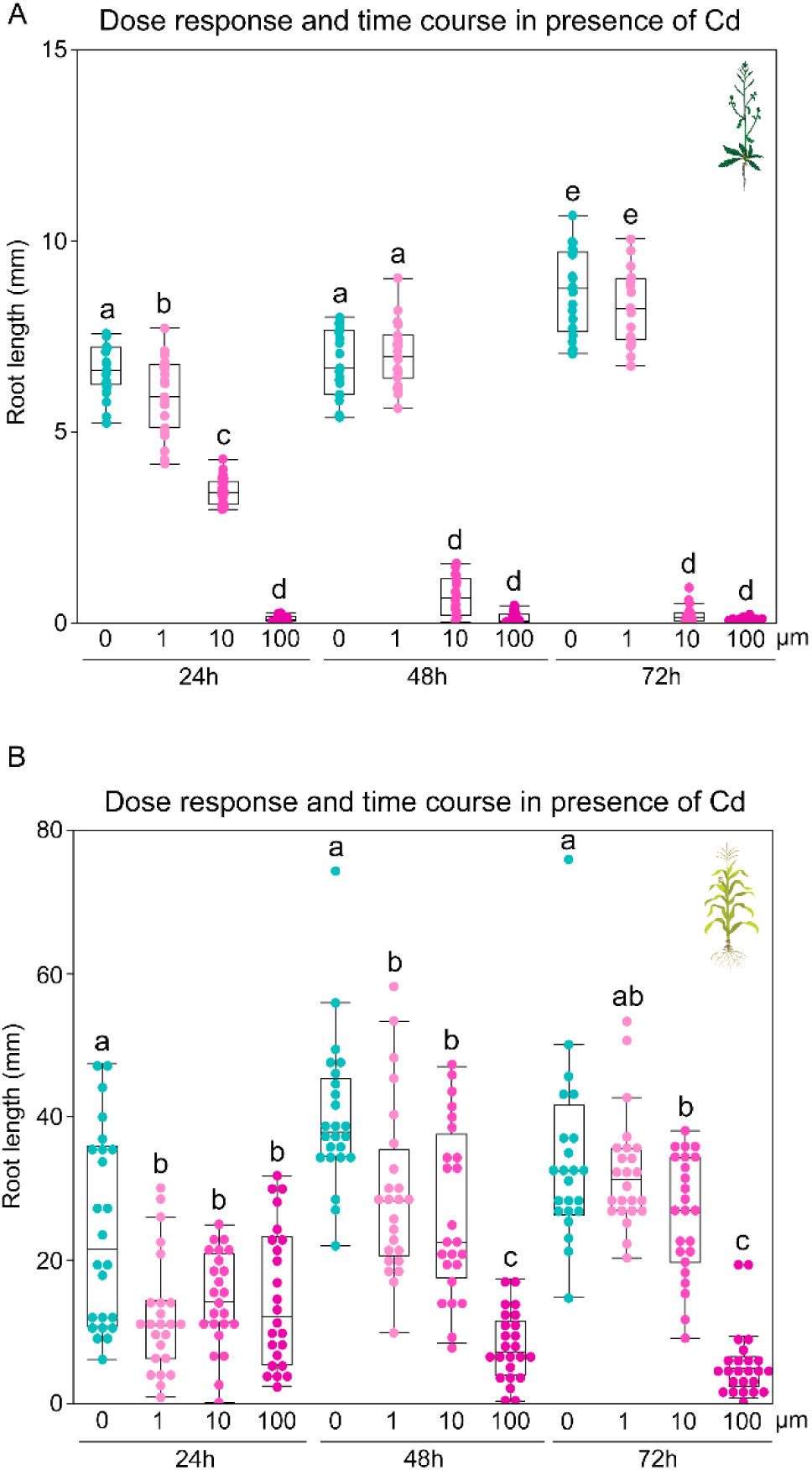
Time course and dose response of Cd in Arabidopsis and maize. (A) Quantification of primary root growth of Arabidopsis after incubating at control, 1µM Cd, 10µM Cd, and 100µM Cd conditions for 24h, 48h, and 72h time points. (B) Quantification of primary root growth of maize after incubating at control, 1µM Cd, 10µM Cd, and 100µM Cd conditions for 24h, 48h, and 72h time points. Statistical test is performed based on Tukey’s Honest test. Groups labeled with the same letter are not statistically different from each other (alpha = 0.05). Boxplots show median values (center line), 25th to 75th interquartile range (box) and 1.5*interquartile range (whiskers).

**Supplemental figure 3.**
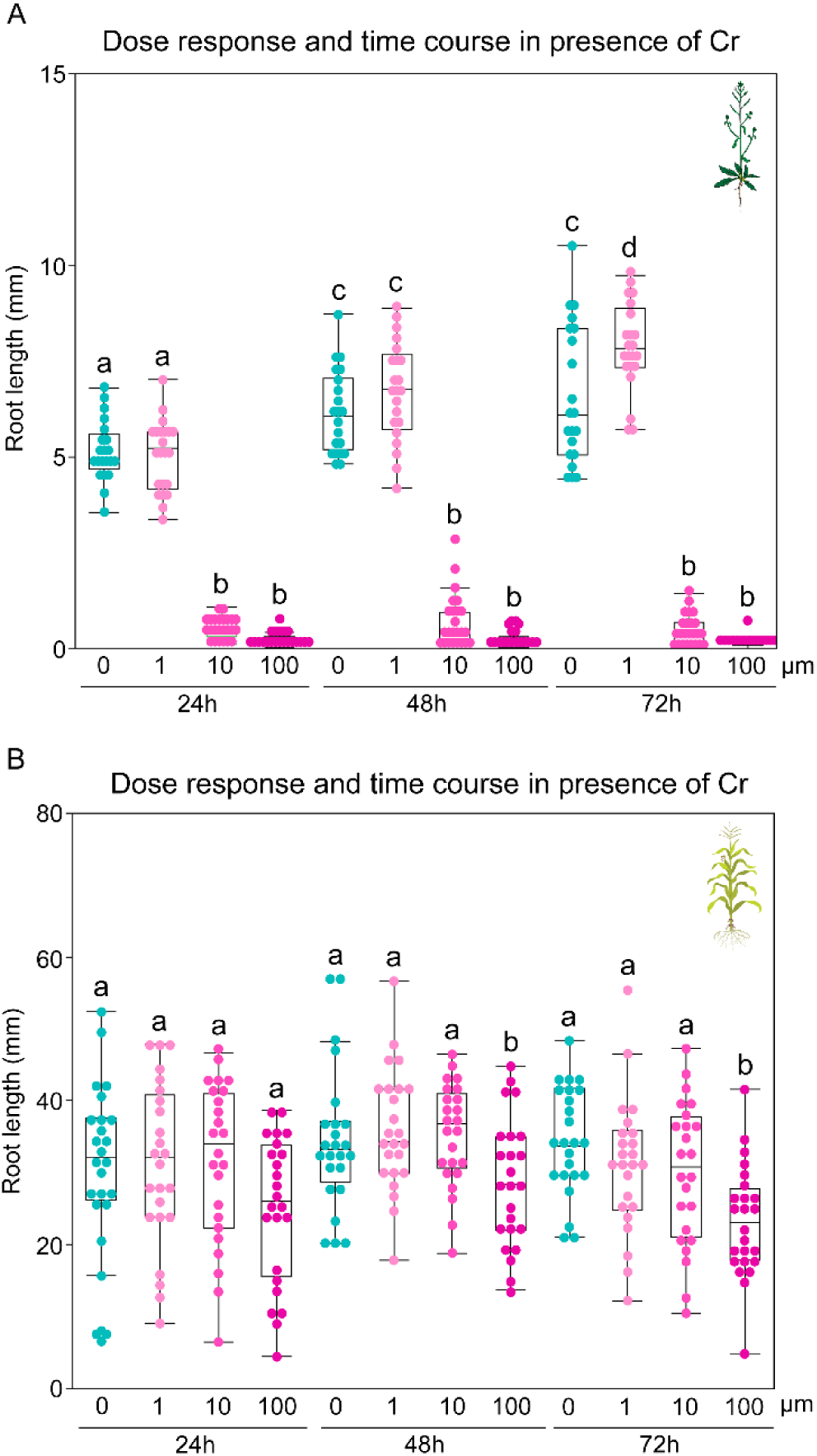
Time course and dose response of Cr in Arabidopsis and maize. (A) Quantification of primary root growth of Arabidopsis after incubating at control, 1µM Cr, 10µM Cr, and 100µM Cr conditions for 24h, 48h, and 72h time points. (B) Quantification of primary root growth of maize after incubating at control, 1µM Cr, 10µM Cr, and 100µM Cr conditions for 24h, 48h, and 72h time points. Statistical test is performed based on Tukey’s Honest test. Groups labeled with the same letter are not statistically different from each other (alpha = 0.05). Boxplots show median values (center line), 25th to 75th interquartile range (box) and 1.5*interquartile range (whiskers).

**Supplemental figure 4.**
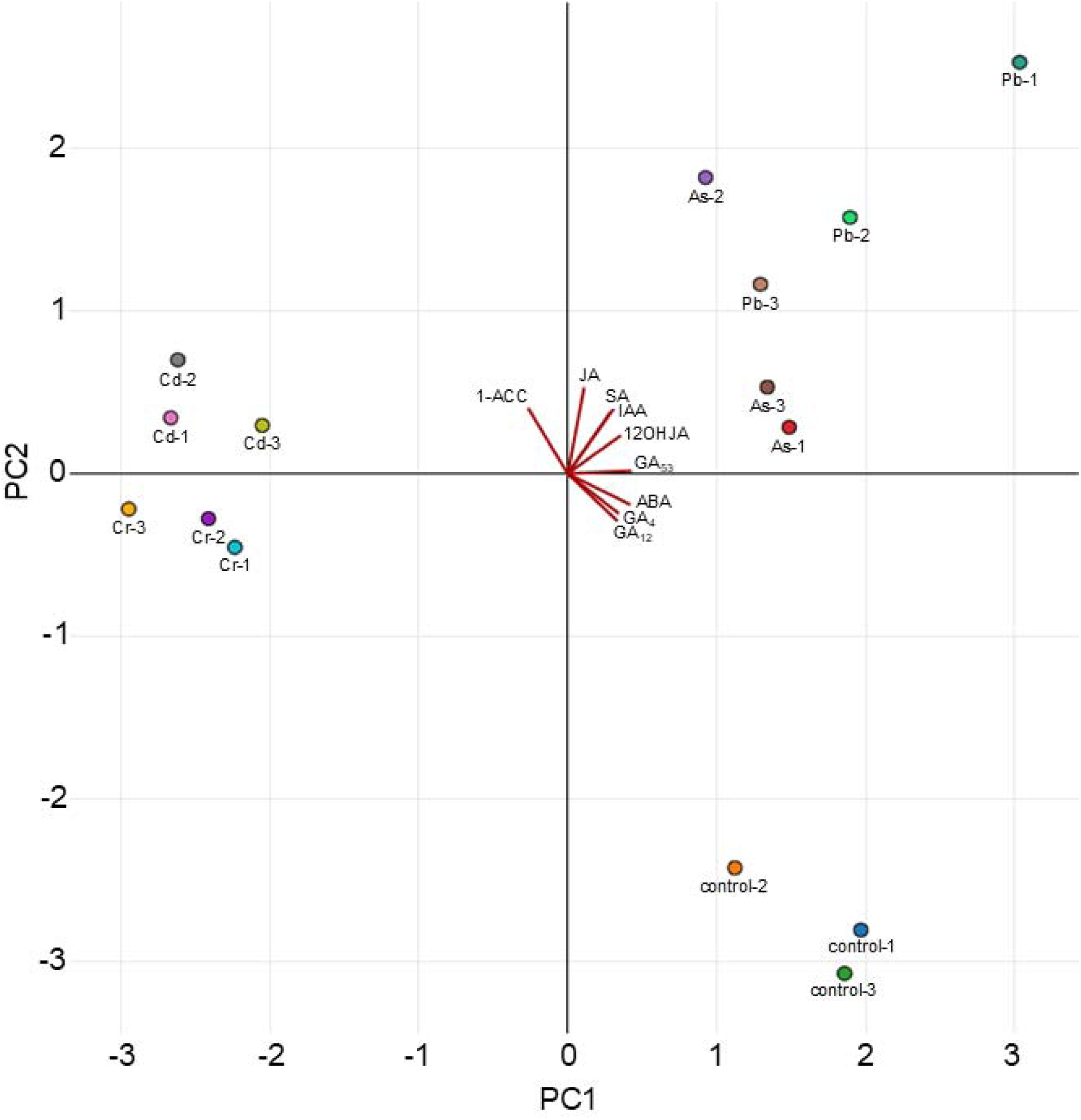
PCA plot for phytohormone quantification in maize root. 3-days-old germinated maize roots were exposed with 10µM As, 10µM Cr, 25µM Cd, and 100µM Pb for 2 days and the root tissues were harvested to measure the phytohormone and generate the PCA plot.

**Supplemental figure 5.**
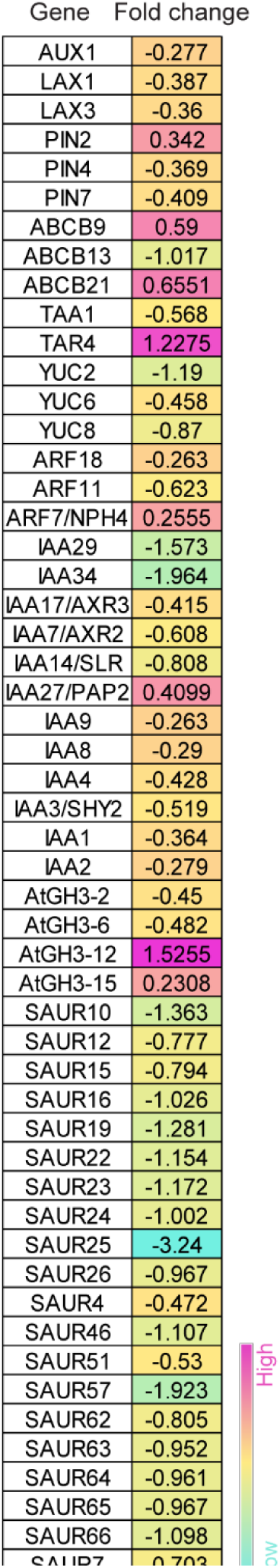
Differentially expressed Arabidopsis genes in presence of lead. Up and down-regulated Arabidopsis Auxin-related genes in presence of 100µM Pb compared to control conditions. Up and down regulation of fold changes are indicated with positive and negative, respectively, values. Gene expression fold change gradient was color coded based on the color gradient scale.

**Supplemental figure 6.**
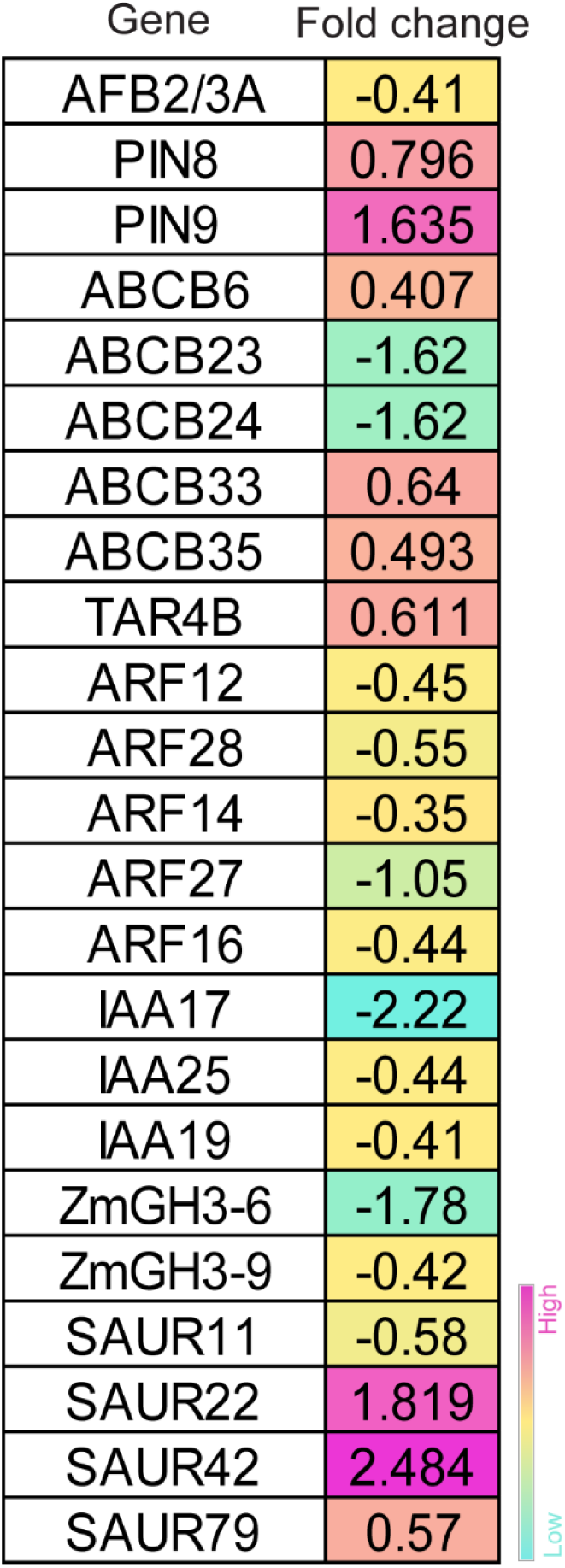
Differentially expressed maize genes in presence of lead. Up and down-regulated Arabidopsis Auxin-related genes in presence of 100µM Pb compared to control conditions. Up and down regulation of fold changes are indicated with positive and negative, respectively, values. Gene expression fold change gradient was color coded based on the color gradient scale.

**Supplemental figure 7.**
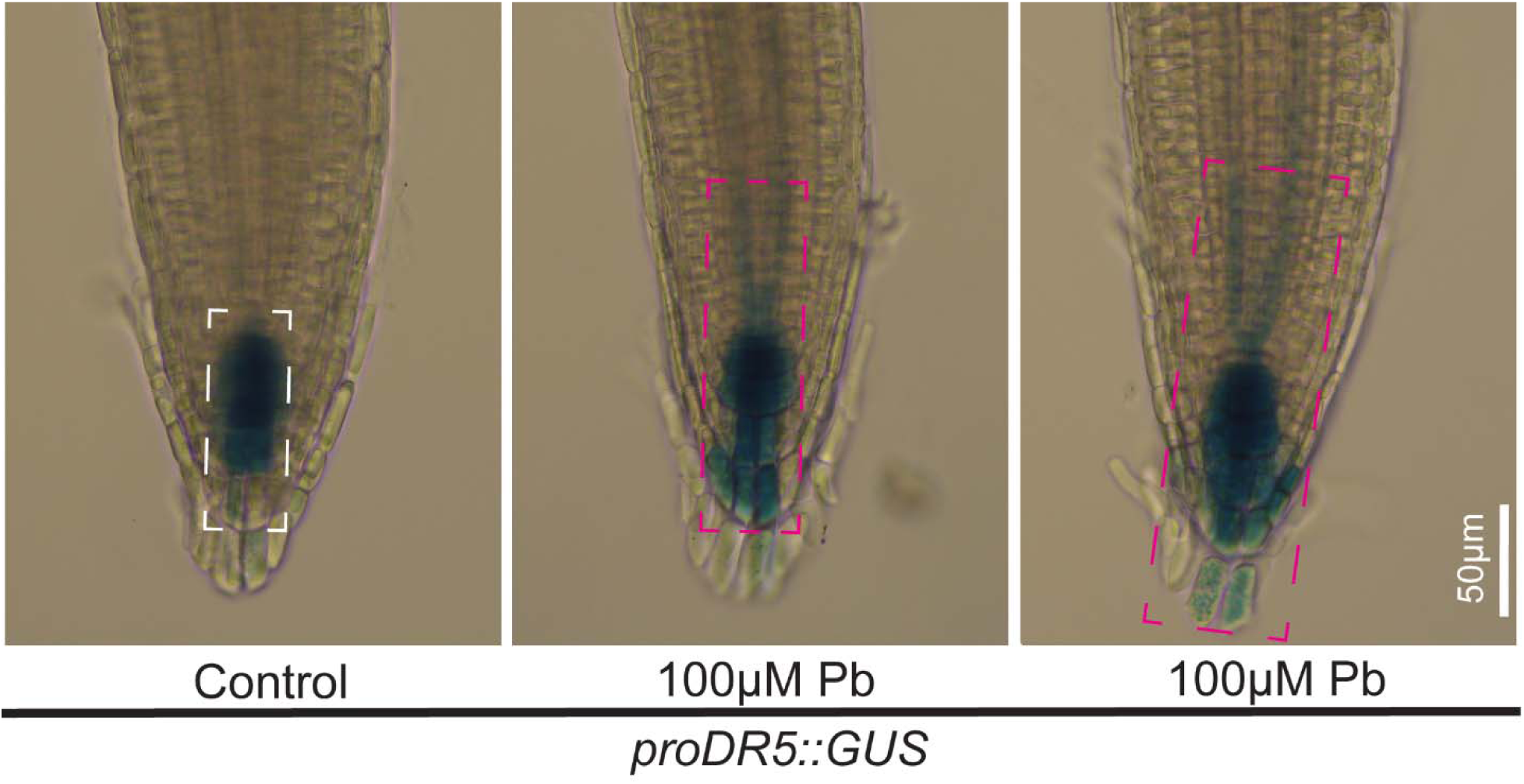
Visualization of proDR5::GUS in presence of lead. proDR5::GUS signal in Arabidopsis root after 2 days of incubation in control and 100µM Pb plates. Scale bar = 50 µm. White and magenta dashed boxes indicate GUS expression regions in control and 100µM Pb-treated roots, respectively.

**Supplemental figure 8.**
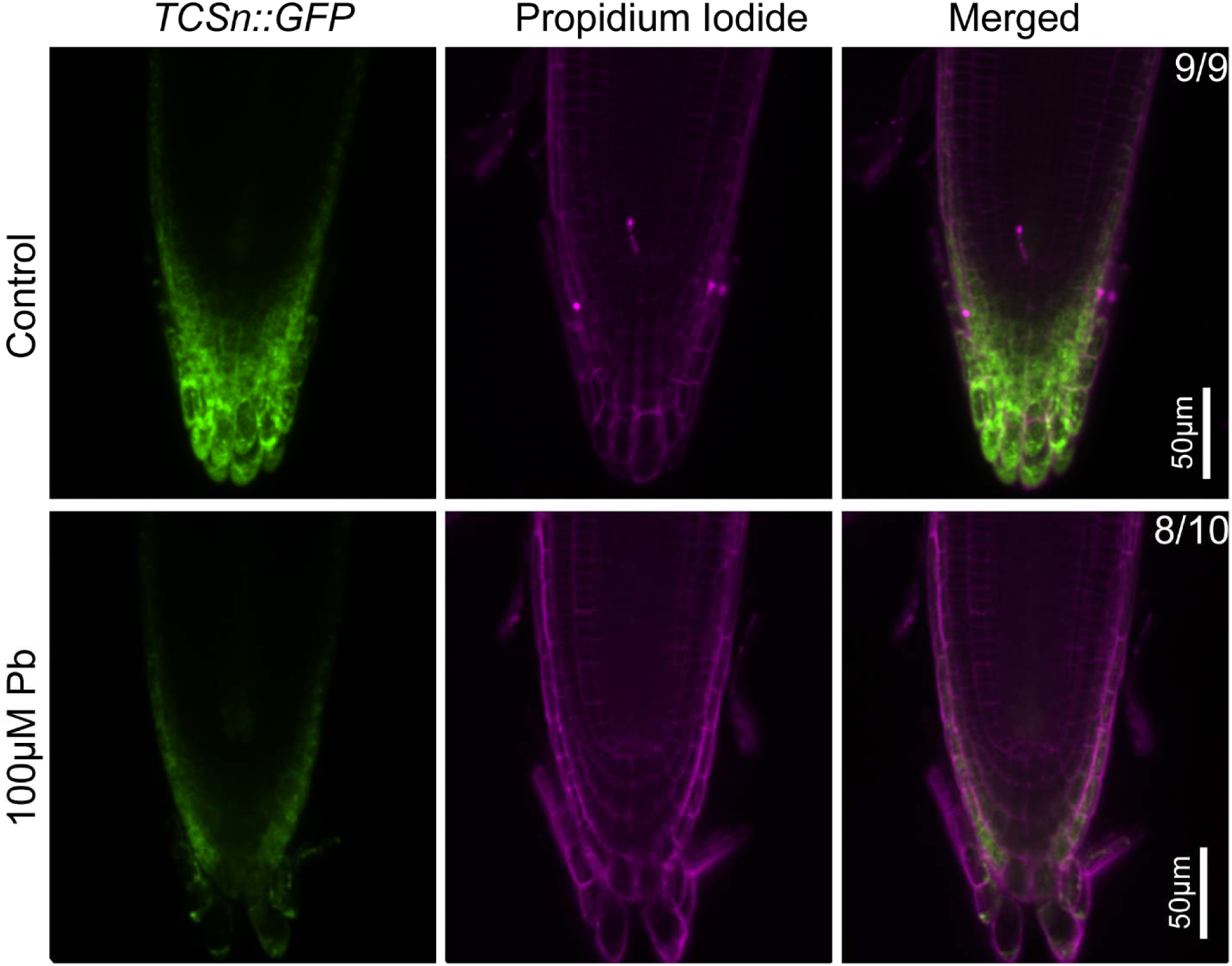
Cytokinin signaling in presence of lead. TCSn::GFP signal in control (upper panel) and 100µM Pb (middle and lower panels) conditions. Roots were counterstained with propidium iodide to visualize individual cells. Scale bar = 50 µm.

**Supplemental figure 9.**
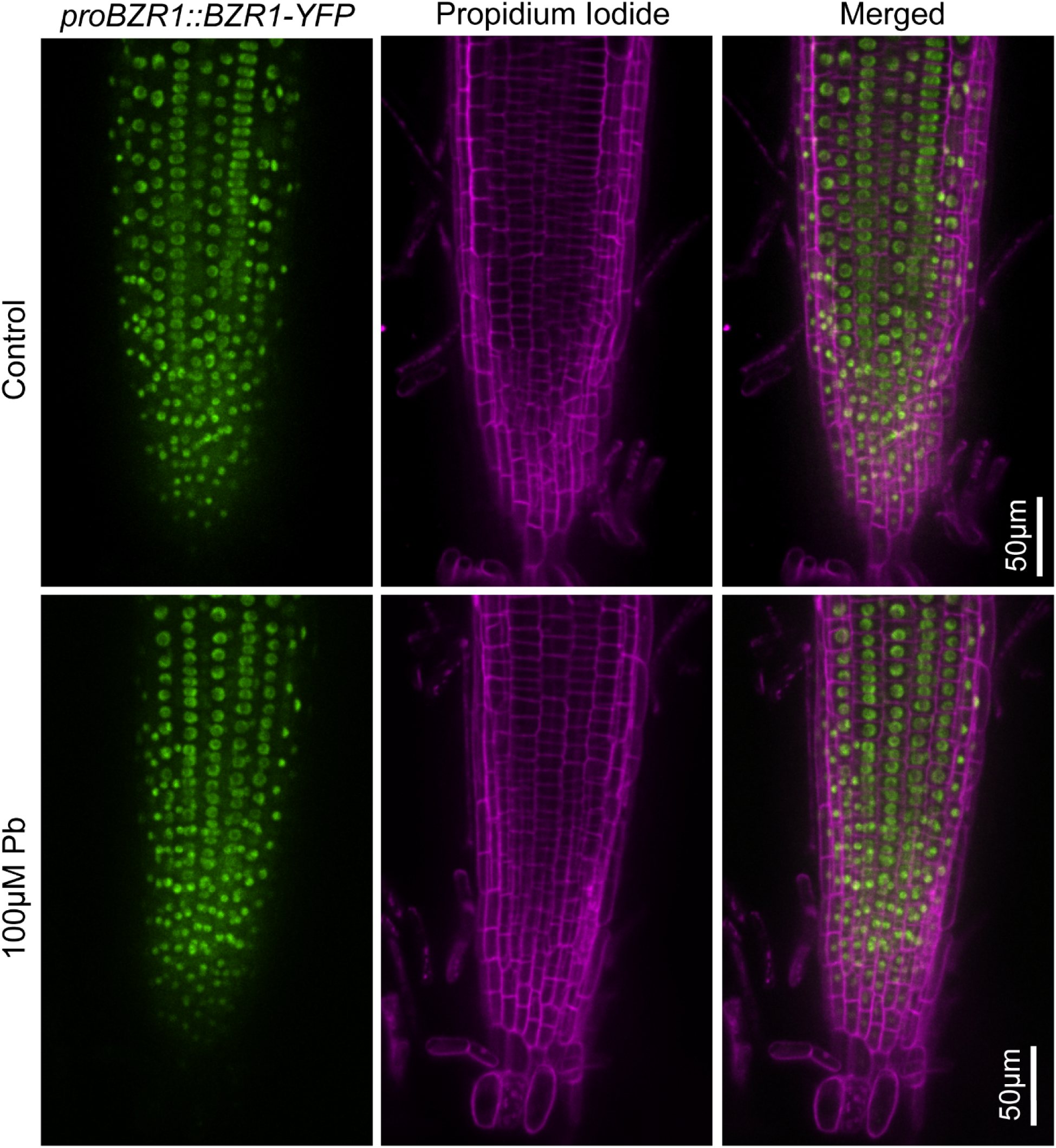
Brassinosteroid signaling in presence of lead. proBZR1::BZR1-GFP signal in control (upper panel) and 100µM Pb (bottom panel) conditions. Roots were counterstained with propidium iodide to visualize individual cells. Scale bar = 50 µm.

**Supplemental figure 10.**
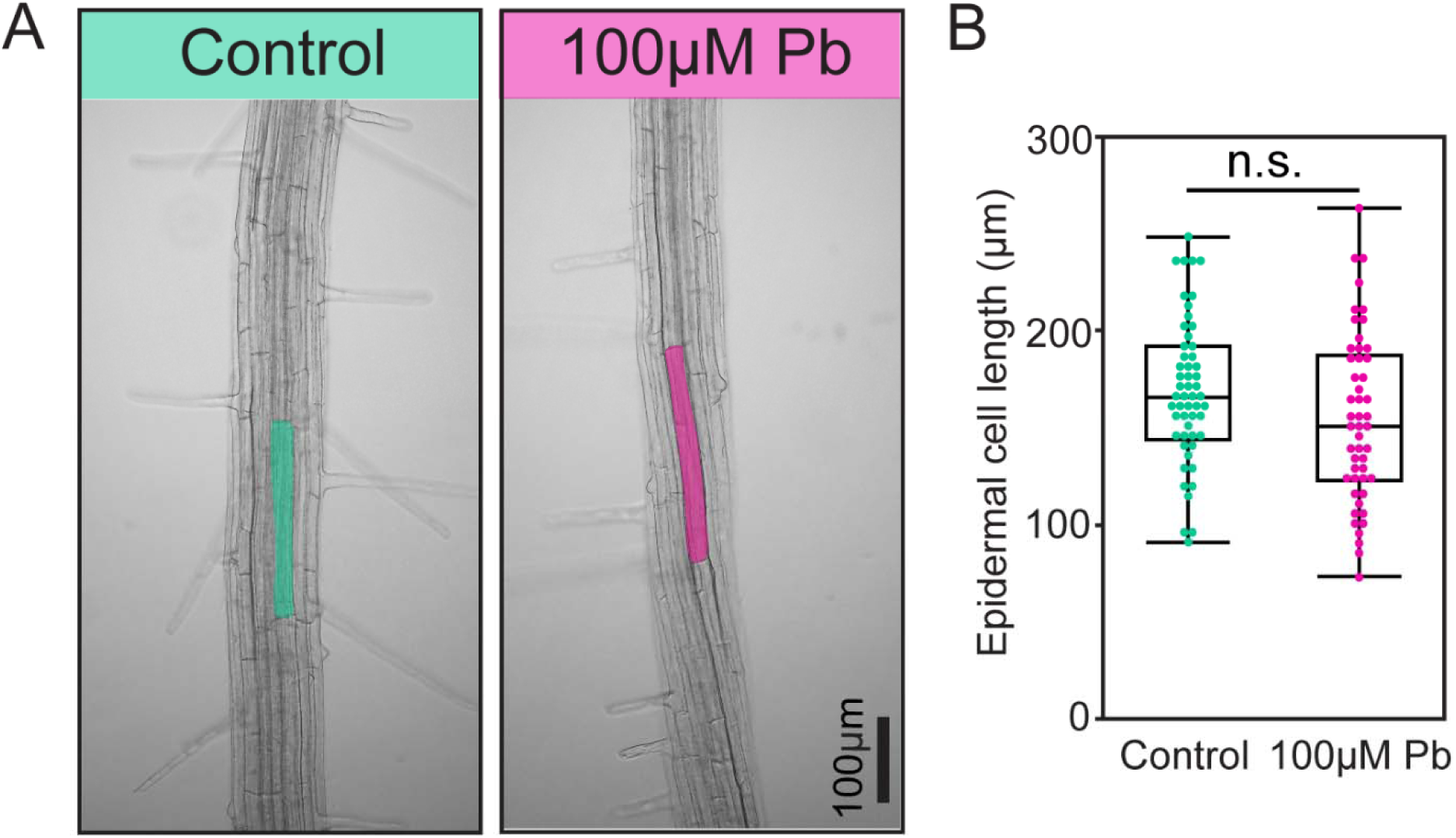
Epidermal cell length in presence of lead. (A) Epidermal cell length was measured after 2 days of incubation in control and 100µM Pb plates. Scale bar = 100µm. (B) Quantification of epidermal cell length from A. Box Plots show median values (center line), 25th to 75th interquartile range (box) and 1.5*interquartile range (whiskers). Student’s t-test was performed for B. Asterisks represent the statistical significance between the means for each treatment. *P < 0.05, **P < 0.01, and ***P < 0.001.

**Supplemental figure 11.**
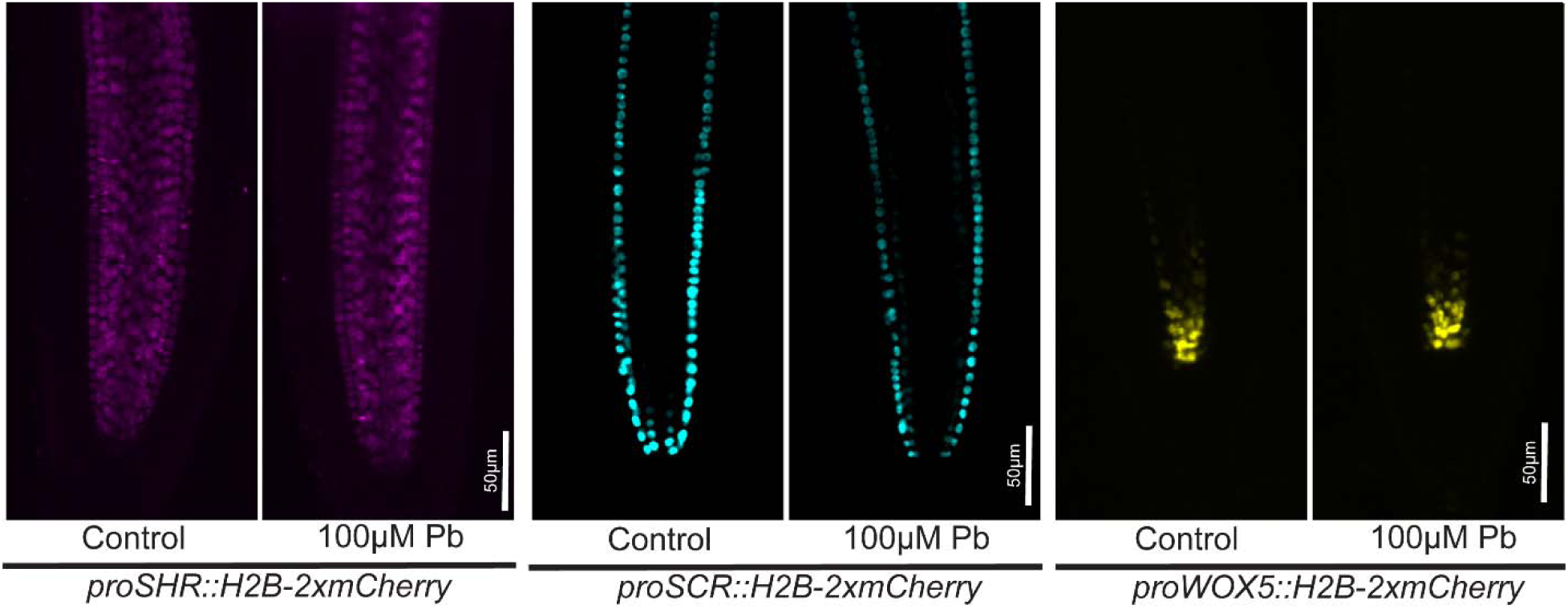
Arabidopsis root developmental marker in presence of lead. proSHR::H2B-2xmCherry, proSCR::H2B-2xmCherry, and proWOX5::H2B-2xmCherry after 2 days of incubation in control and 100µM Pb plates. Scale bar = 50µm.

**Supplemental figure 12.**
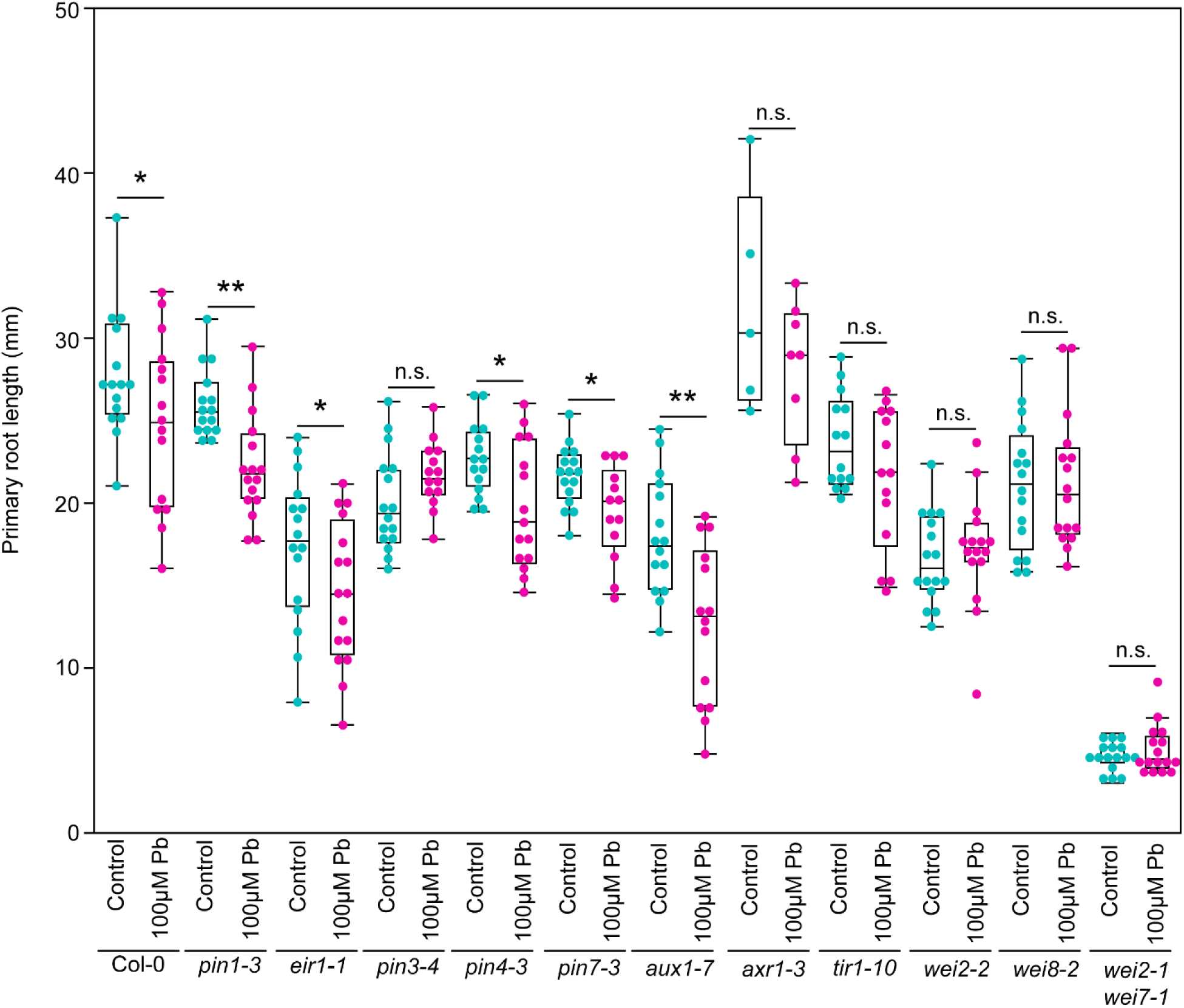
Screening of auxin mutants in presence of lead. Auxin transport (*pin1-3*, *eir1-1*, *pin3-4*, *pin4-3*, *pin7-3*, and *aux1-7*), signaling (*tir1-10* and *axr1-3*), and biosynthesis (*wei2-2*, *wei8-2*, and *wei2-1 wei7-1*) mutants were transferred to control and 100µM Pb to observe the primary root growth response. Student’s t-test was performed. Asterisks represent the statistical significance between the means for each treatment. *P < 0.05, **P < 0.01, and ***P < 0.001.

**Supplemental Table 1.** Quantification of brassinosteroid hormones in maize root.

**Supplemental Table 2.** Sequencing information and statistics.

## Notes

### Competing Interest Statement

The authors have declared no competing interest.

### Summary of Updates

The manuscript was revised significantly, including time course and dose response for all different heavy metals in Arabidopsis and maize. Additionally, we performed the reverse genetic screening with auxin mutants and quantitative cell biology for auxin reporters.

