## Supplemental figures for "Toxic metal, Lead (Pb) alters columella-specific auxin distribution inhibiting root growth and gravitropic response in Arabidopsis and maize"

**Supplemental figure 1**

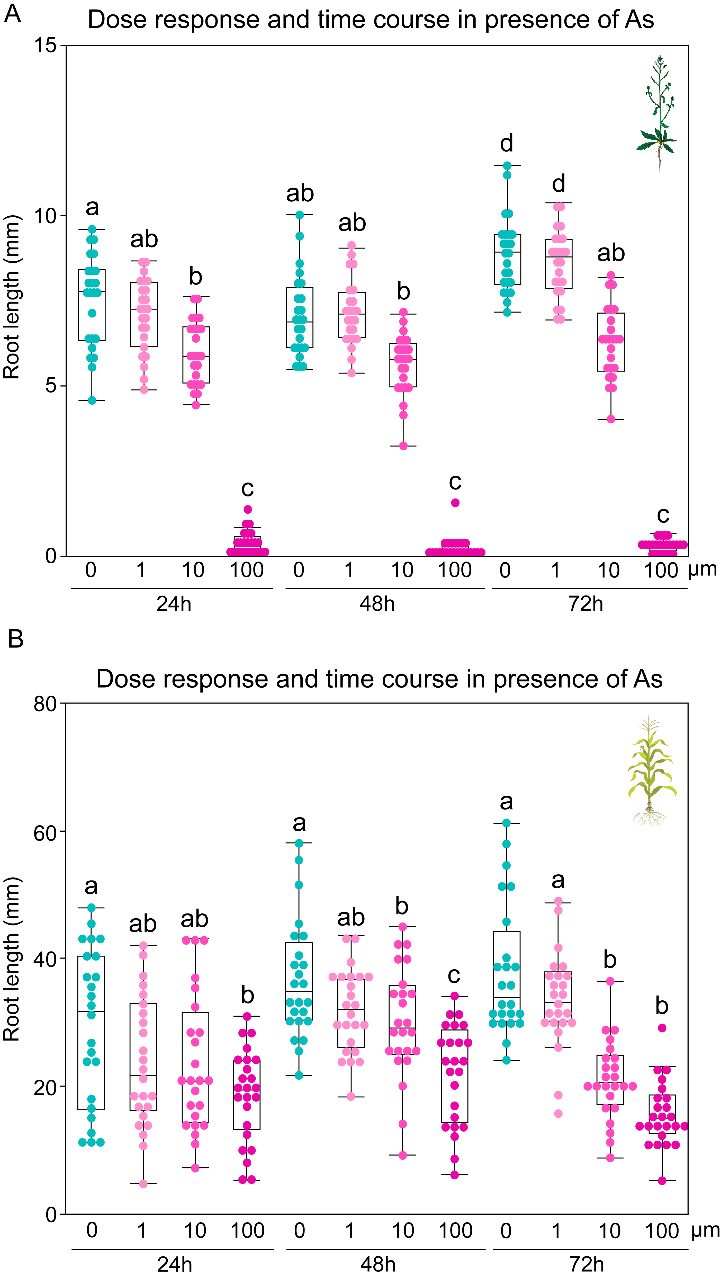

**Supplemental figure 1.** Time course and dose response of As in Arabidopsis and maize.

(A) Quantification of primary root growth of Arabidopsis after incubating at control, 1µM As, 10µM As, and 100µM As conditions for 24h, 48h, and 72h time points. (B) Quantification of primary root growth of maize after incubating at control, 1µM As, 10µM As, and 100µM As conditions for 24h, 48h, and 72h time points. Statistical test is performed based on Tukey's Honest test. Groups labeled with the same letter are not statistically different from each other (alpha = 0.05). Boxplots show median values (center line), 25th to 75th interquartile range (box) and 1.5*interquartile range (whiskers).

**Supplemental figure 2**

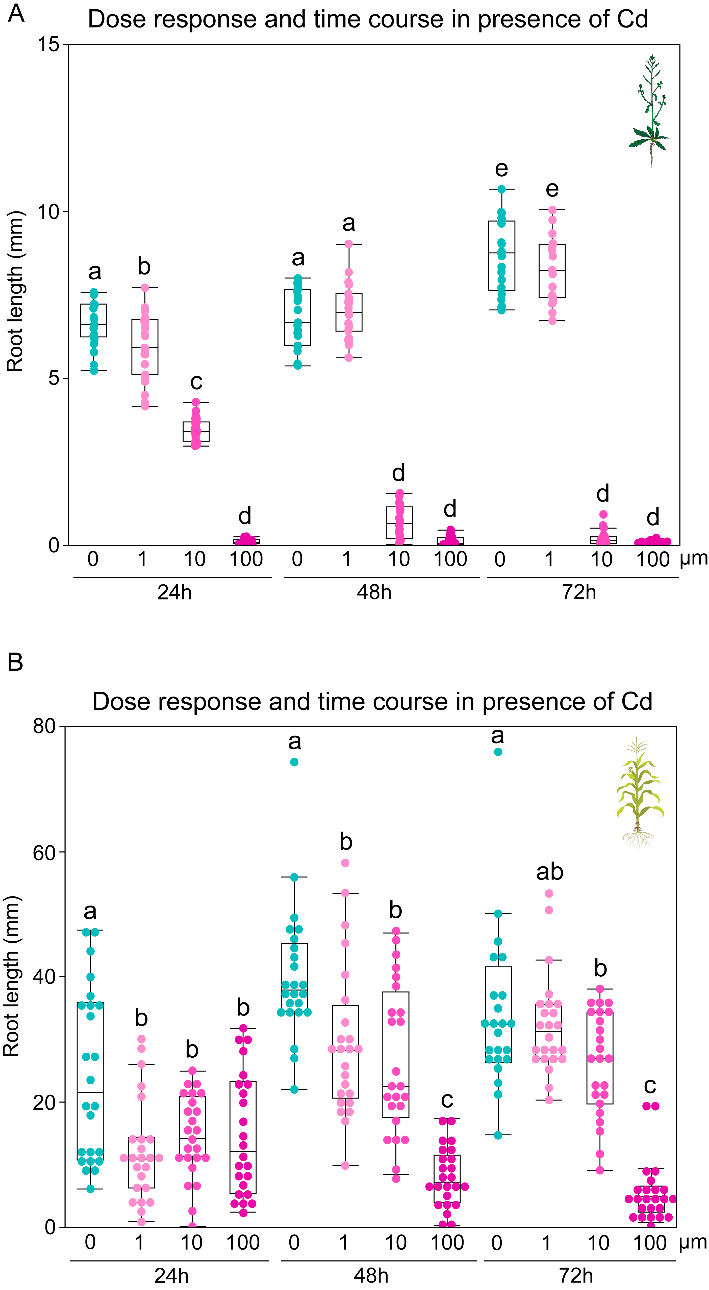

**Supplemental figure 2.** Time course and dose response of Cd in Arabidopsis and maize.

(A) Quantification of primary root growth of Arabidopsis after incubating at control, 1µM Cd, 10µM Cd, and 100µM Cd conditions for 24h, 48h, and 72h time points. (B) Quantification of primary root growth of maize after incubating at control, 1µM Cd, 10µM Cd, and 100µM Cd conditions for 24h, 48h, and 72h time points. Statistical test is performed based on Tukey's Honest test. Groups labeled with the same letter are not statistically different from each other (alpha = 0.05). Boxplots show median values (center line), 25th to 75th interquartile range (box) and 1.5*interquartile range (whiskers).

**Supplemental figure 3**

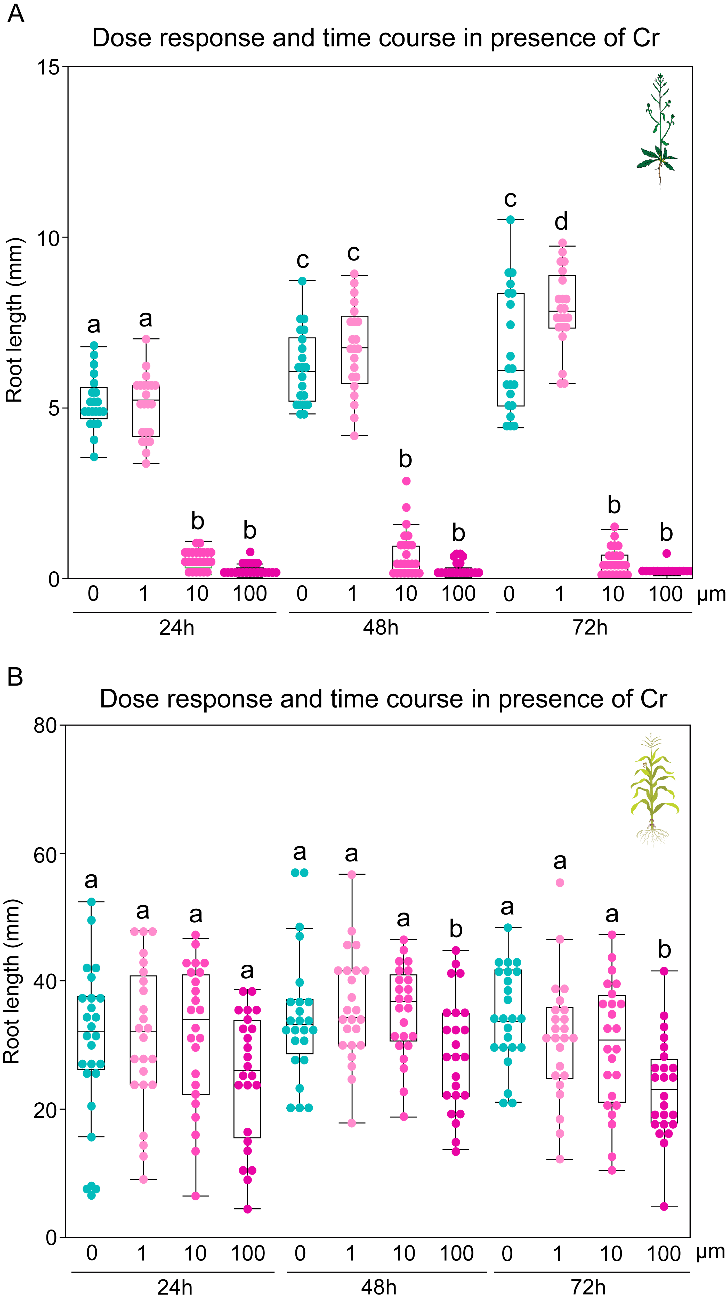

**Supplemental figure 3.** Time course and dose response of Cr in Arabidopsis and maize.

(A) Quantification of primary root growth of Arabidopsis after incubating at control, 1µM Cr, 10µM Cr, and 100µM Cr conditions for 24h, 48h, and 72h time points. (B) Quantification of primary root growth of maize after incubating at control, 1µM Cr, 10µM Cr, and 100µM Cr conditions for 24h, 48h, and 72h time points. Statistical test is performed based on Tukey's Honest test. Groups labeled with the same letter are not statistically different from each other (alpha = 0.05). Boxplots show median values (center line), 25th to 75th interquartile range (box) and 1.5*interquartile range (whiskers).

**Supplemental figure 4**

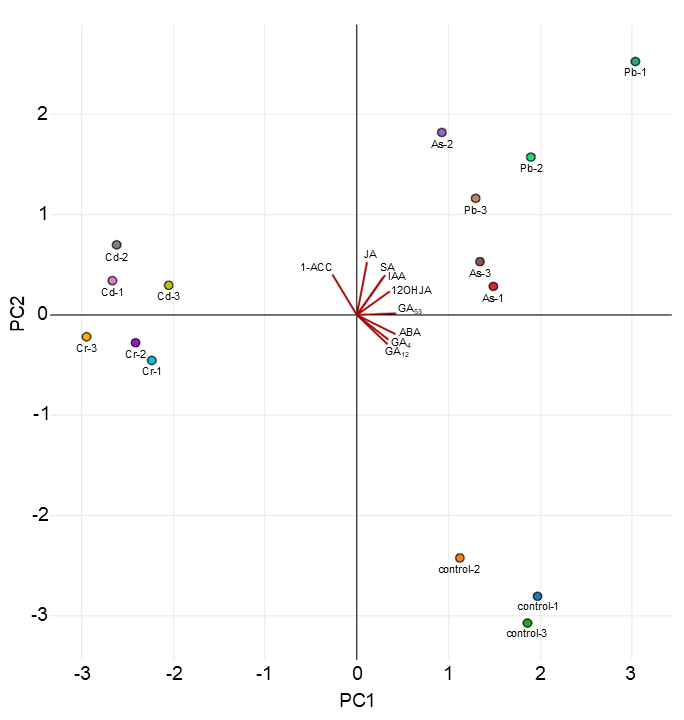

**Supplemental figure 4.** PCA plot for phytohormone quantification in maize root.

3-days-old germinated maize roots were exposed with 10µM As, 10µM Cr, 25µM Cd, and 100µM Pb for 2 days and the root tissues were harvested to measure the phytohormone and generate the PCA plot.

**Supplemental figure 5**

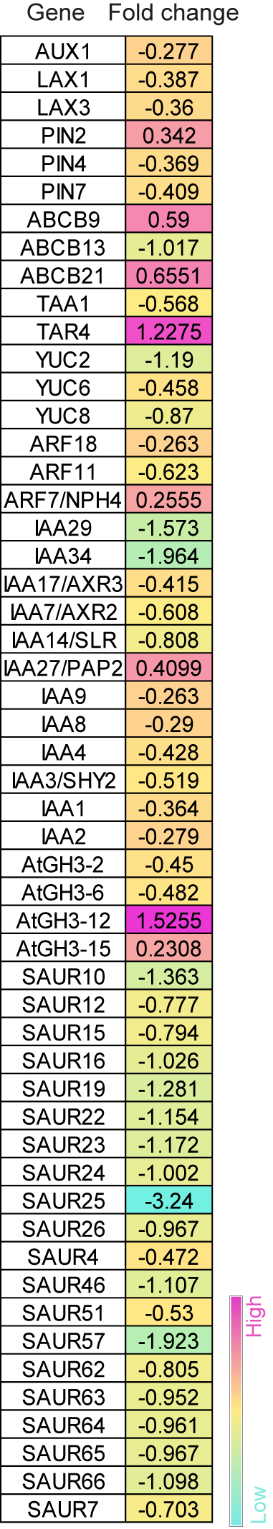

**Supplemental figure 5.** Differentially expressed Arabidopsis genes in presence of lead. Up and down-regulated Arabidopsis Auxin-related genes in presence of 100µM Pb compared to control conditions. Up and down regulation of fold changes are indicated with positive and negative, respectively, values. Gene expression fold change gradient was color coded based on the color gradient scale.

**Supplemental figure 6**

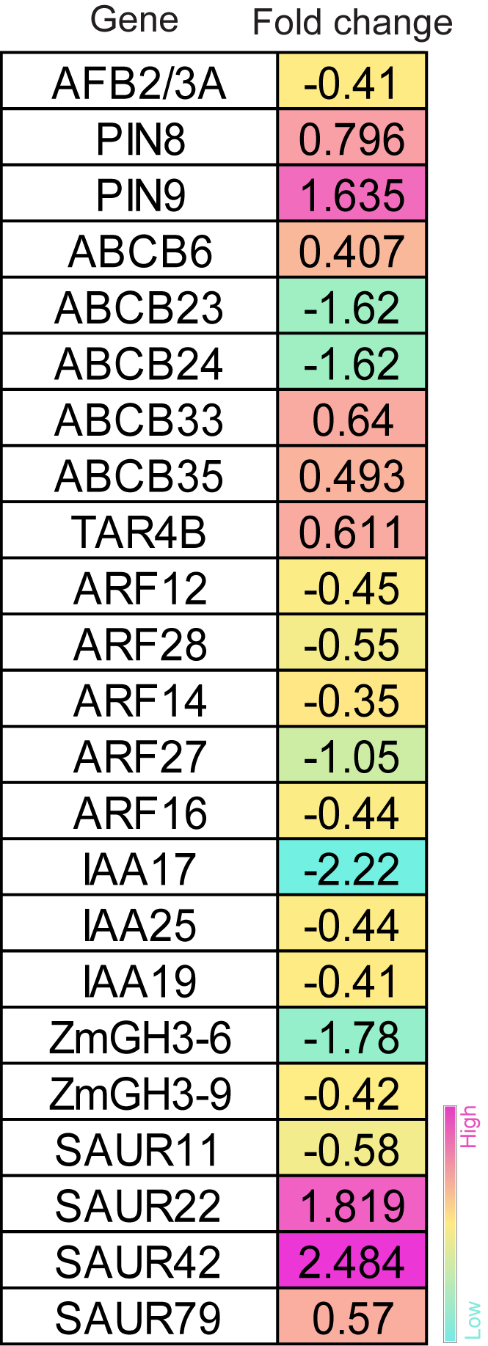

**Supplemental figure 6.** Differentially expressed maize genes in presence of lead. Up and down-regulated Arabidopsis Auxin-related genes in presence of 100µM Pb compared to control conditions. Up and down regulation of fold changes are indicated with positive and negative, respectively, values. Gene expression fold change gradient was color coded based on the color gradient scale.

**Supplemental figure 7**

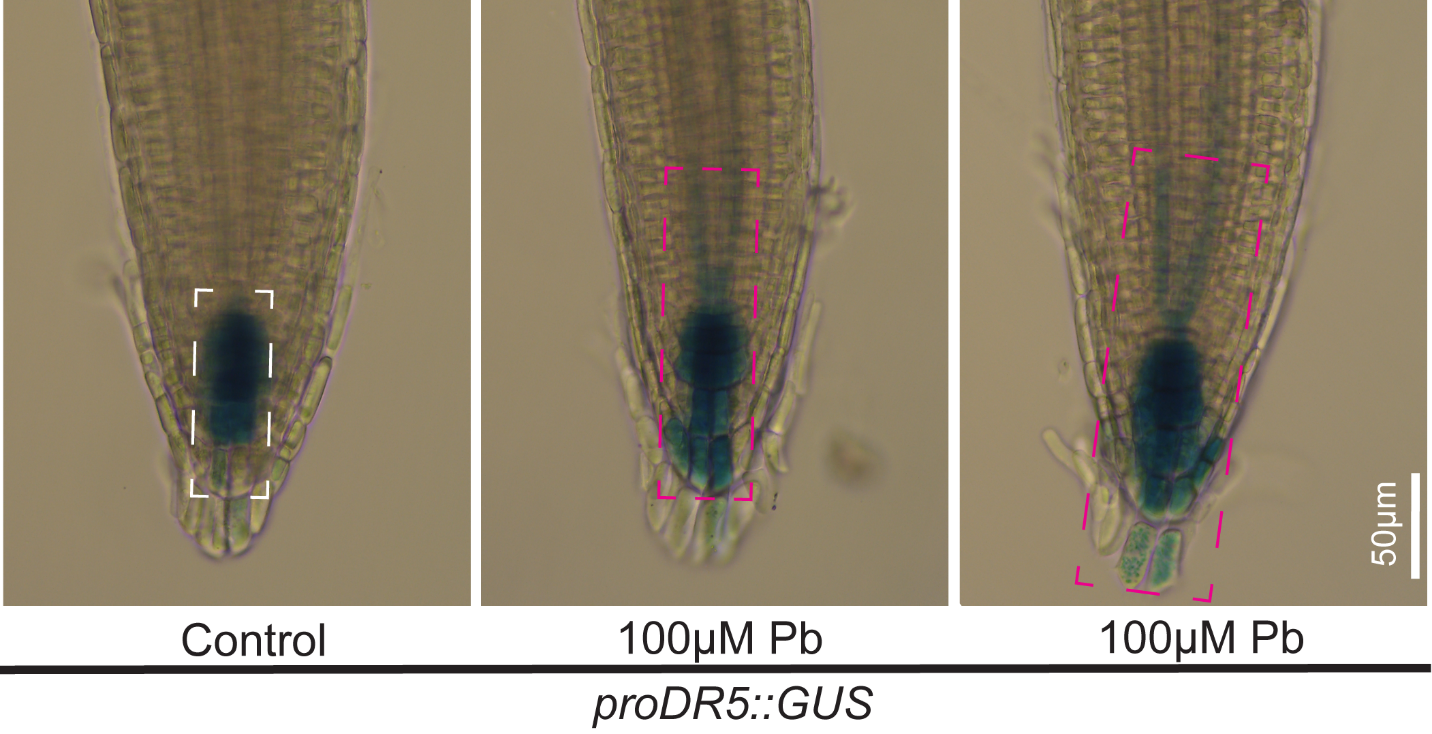

**Supplemental figure 7.** Visualization of proDR5::GUS in presence of lead.

proDR5::GUS signal in Arabidopsis root after 2 days of incubation in control and 100µM Pb plates. Scale bar = 50 µm. White and magenta dashed boxes indicate GUS expression regions in control and 100µM Pb-treated roots, respectively.

**Supplemental figure 8**

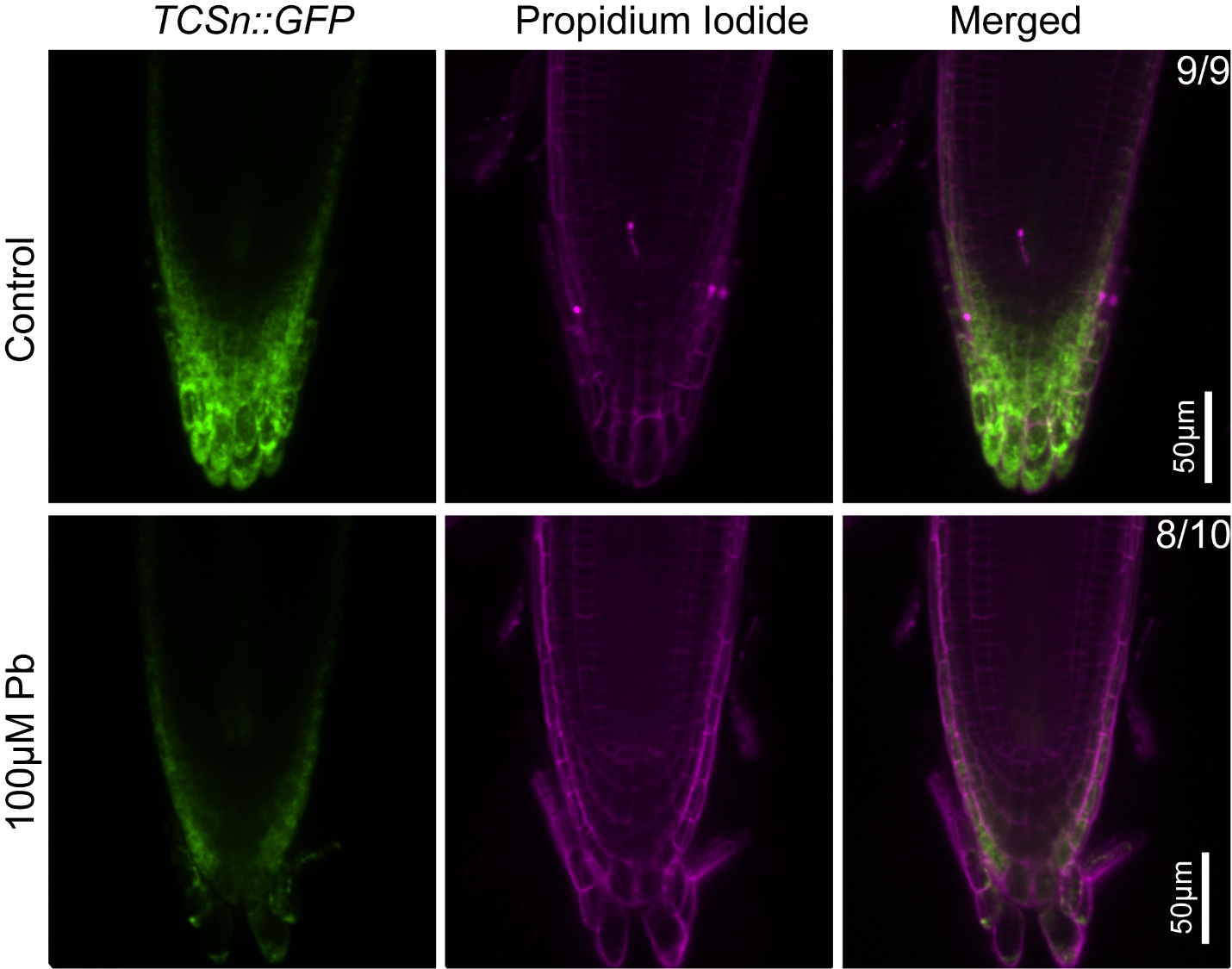

**Supplemental figure 8.** Cytokinin signaling in presence of lead.

TCSn::GFP signal in control (upper panel) and 100µM Pb (middle and lower panels) conditions. Roots were counterstained with propidium iodide to visualize individual cells. Scale bar = 50 µm.

**Supplemental figure 9**

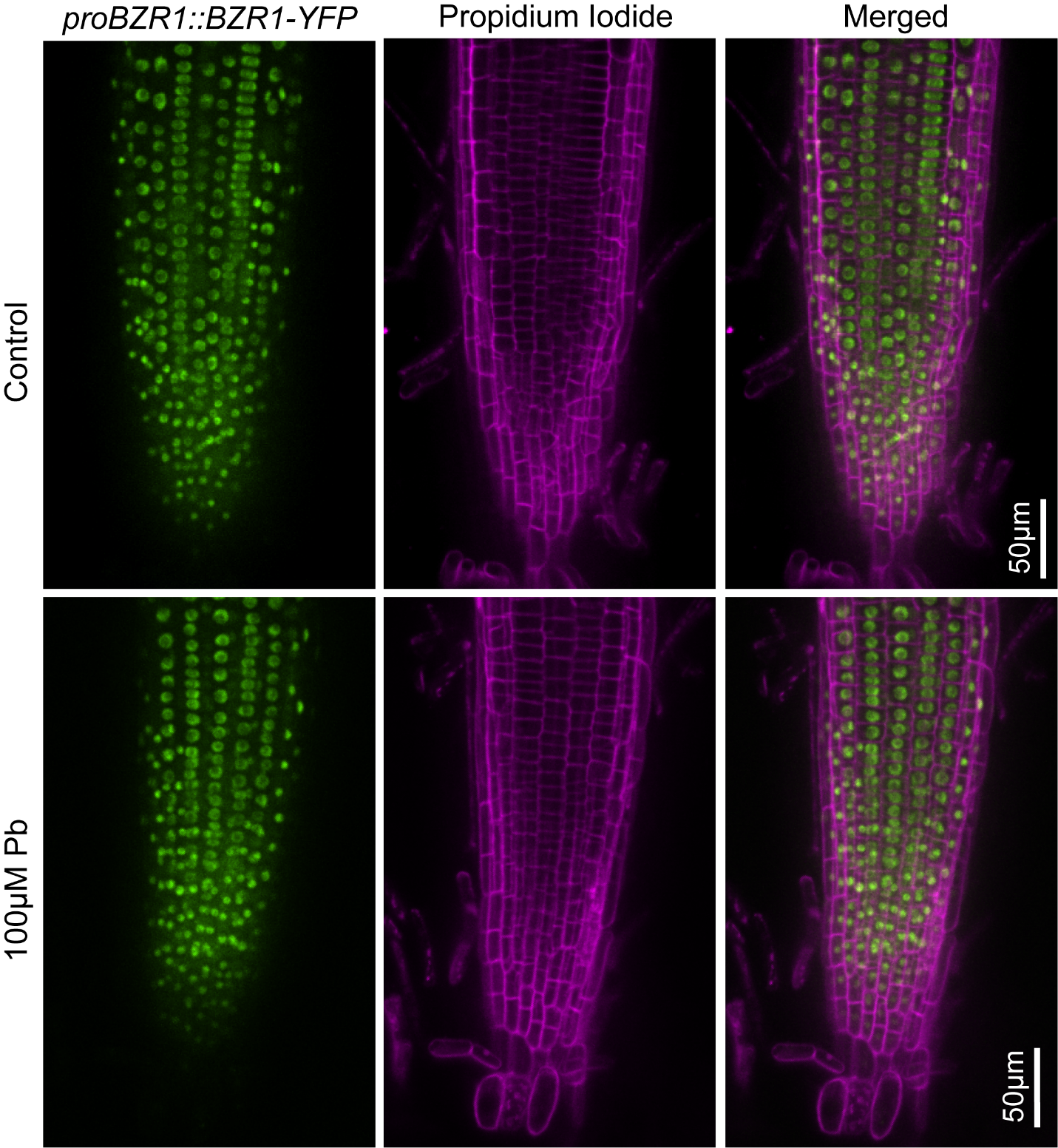

**Supplemental figure 9.** Brassinosteroid signaling in presence of lead.

proBZR1::BZR1-GFP signal in control (upper panel) and 100µM Pb (bottom panel) conditions. Roots were counterstained with propidium iodide to visualize individual cells. Scale bar = 50 µm.

**Supplemental figure 10**

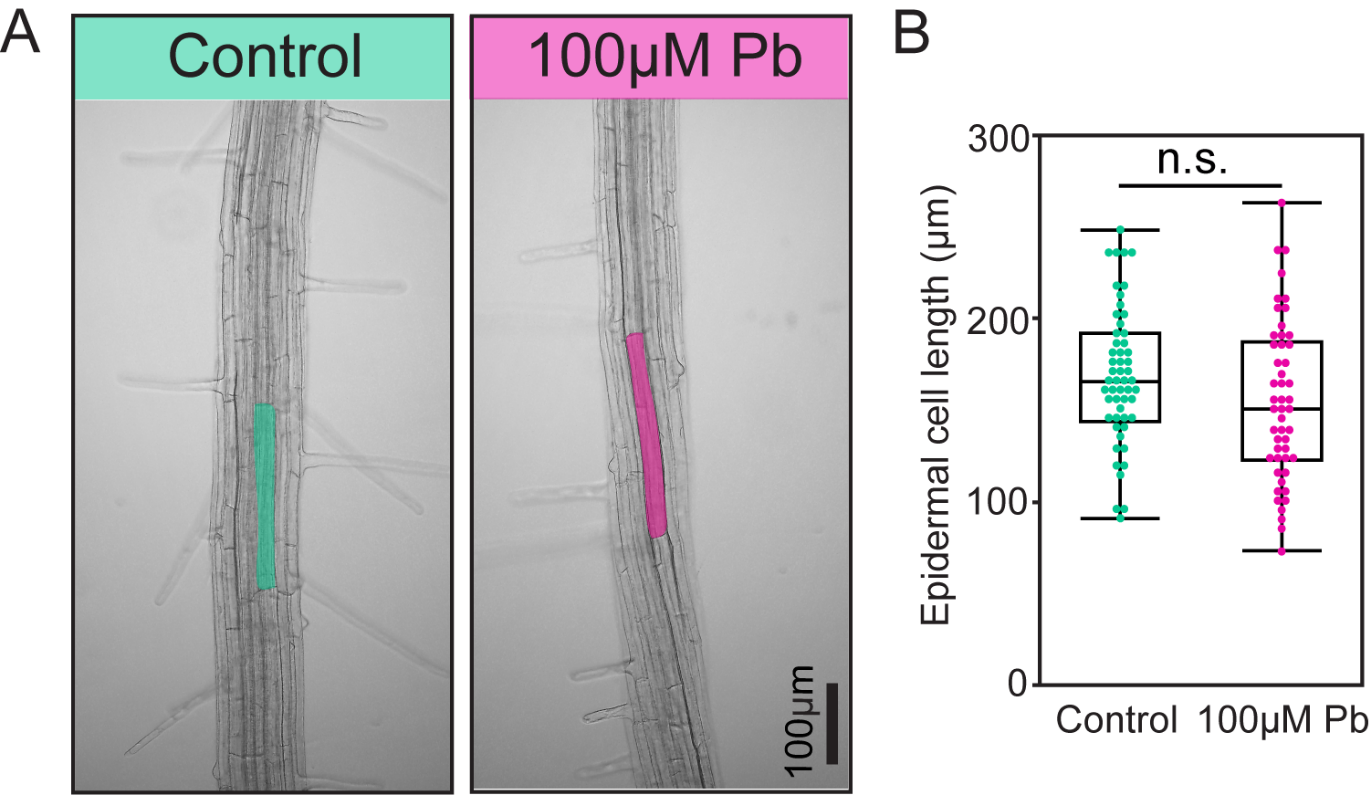

**Supplemental figure 10.** Epidermal cell length in presence of lead.

(A) Epidermal cell length was measured after 2 days of incubation in control and 100µM Pb plates. Scale bar = 100µm. (B) Quantification of epidermal cell length from A. Box Plots show median values (center line), 25th to 75th interquartile range (box) and 1.5*interquartile range (whiskers). Student’s t-test was performed for B. Asterisks represent the statistical significance between the means for each treatment. *P < 0.05, **P < 0.01, and ***P < 0.001.

**Supplemental figure 11**

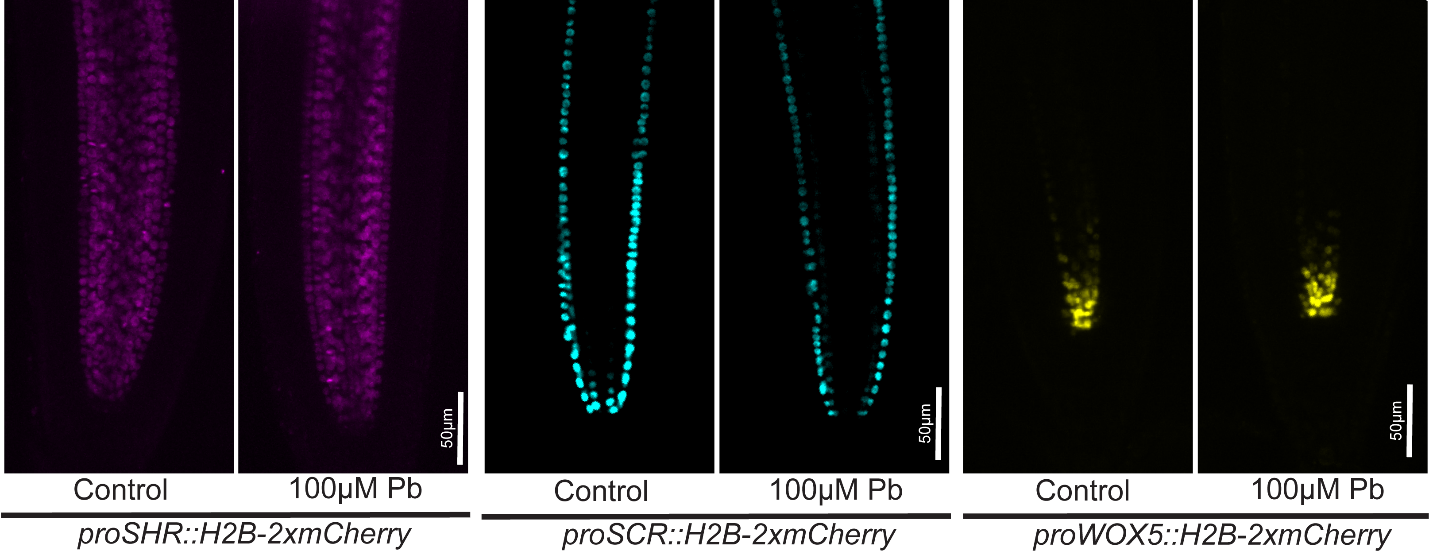

**Supplemental figure 11.** Arabidopsis root developmental marker in presence of lead.

proSHR::H2B-2xmCherry, proSCR::H2B-2xmCherry, and proWOX5::H2B-2xmCherry after 2 days of incubation in control and 100µM Pb plates. Scale bar = 50µm.

**Supplemental figure 12**

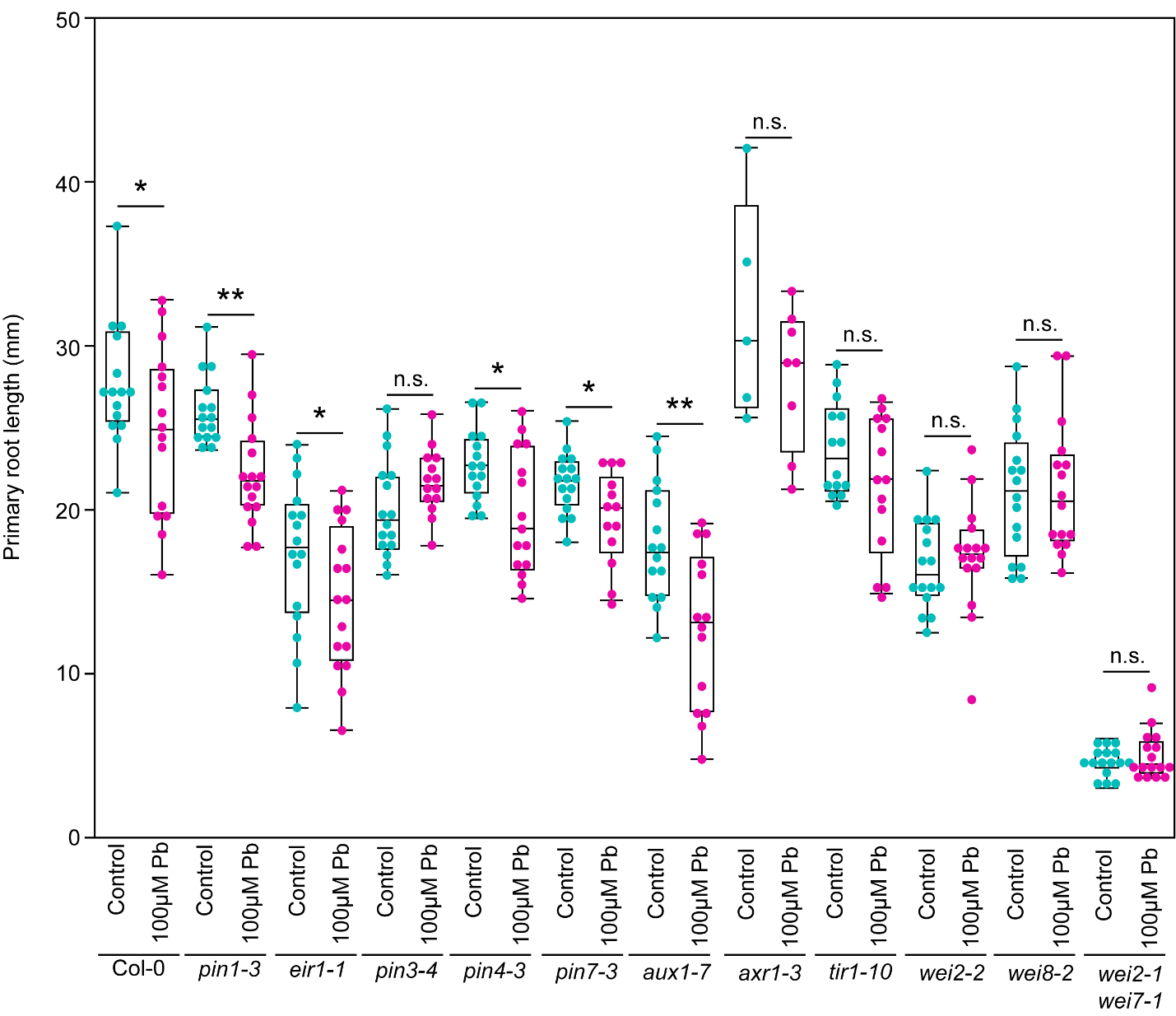

**Supplemental figure 12.** Screening of auxin mutants in presence of lead.

Auxin transport (*pin1-3*, *eir1-1*, *pin3-4*, *pin4-3*, *pin7-3*, and *aux1-7*), signaling (*tir1-10* and *axr1-3*), and biosynthesis (*wei2-2*, *wei8-2*, and *wei2-1 wei7-1*) mutants were transferred to control and 100µM Pb to observe the primary root growth response. Student’s t-test was performed. Asterisks represent the statistical significance between the means for each treatment. *P < 0.05, **P < 0.01, and ***P < 0.001.
