## Supplemental Table 1 for "Toxic metal, Lead (Pb) alters columella-specific auxin distribution inhibiting root growth and gravitropic response in Arabidopsis and maize"

Supplemental Table 1: Steroid hormone quantification from maize root during control, As, Cd, Cr, and Pb conditions.

| **Content (ng/g FW)** | **Control** | **10µM Cr** | **25µM Cd** | **100µM Pb** | **10µM As** |
| --- | --- | --- | --- | --- | --- |
| **Isofucosterol** | 313.13 ± 30.55 | 261.26 ± 11.61 | 342.65 ± 93.25 | 310.34 ± 66.63 | 301.27 ± 87.68 |
| **Stigmasterol** | 1217.11 ± 178.41 | 1095.34 ± 58.05 | 1170.72 ± 165.10 | 1302.59 ± 115.21 | 1374.91 ± 74.78 |
| **Sitosterol** | 3616.56 ± 1335.10 | 3002.57 ± 2103.77 | 3768.93 ± 1351.74 | 5022.08 ± 1579.88 | 3843.52 ± 1037.71 |
| **24-methylenecholesterol** | 181.05 ± 38.53 | 161.51 ± 48.33 | 202.19 ± 47.80 | 185.77 ± 57.74 | 140.94 ± 40.60 |
| **Campesterol** | 2003.32 ± 176.67 | 1982.69 ± 27.47 | 2004.55 ± 266.61 | 2127.82 ± 109.63 | 2192.31 ± 169.72 |
| **Campestanol** | 65.45 ± 3.58 | 64.68 ± 20.36 | 76.60 ± 18.55 | 80.14 ± 16.1 | 75.38 ± 14.37 |
| **6-oxocampestanol** | 125.36 ± 33.33 | 140.38 ± 36.36 | 121.38 ± 33.05 | 169.07 ± 60.38 | 153.80 ± 56.09 |
