## Supplemental Table 2 for "Toxic metal, Lead (Pb) alters columella-specific auxin distribution inhibiting root growth and gravitropic response in Arabidopsis and maize"

Supplemental Table 1. Sequencing information and statistics.

| Sample Accession Number | BioProject Number | Species and Sample Type | Treatment | File Name ID | Number of Paired-end Reads | Alignment rate |
| --- | --- | --- | --- | --- | --- | --- |
| SAMN48150518 | PRJNA1255621 | Maize RNA | Control | Zm1 | 32805461 | 89.88% |
| SAMN48150519 | PRJNA1255621 | Maize RNA | Control | Zm2 | 28309098 | 89.49% |
| SAMN48150520 | PRJNA1255621 | Maize RNA | Control | Zm3 | 32343118 | 89.07% |
| SAMN48150521 | PRJNA1255621 | Maize RNA | Lead | Zm4 | 38100741 | 86.57% |
| SAMN48150522 | PRJNA1255621 | Maize RNA | Lead | Zm5 | 35723413 | 89.00% |
| SAMN48150523 | PRJNA1255621 | Maize RNA | Lead | Zm6 | 36459353 | 89.76% |
| SAMN48150797 | PRJNA1255621 | Arabidopsis RNA | Control | Control-1 | 29109904 | 96.74% |
| SAMN48150798 | PRJNA1255621 | Arabidopsis RNA | Control | Control-2 | 24007962 | 96.29% |
| SAMN48150799 | PRJNA1255621 | Arabidopsis RNA | Control | Control-3 | 21156599 | 96.14% |
| SAMN48150800 | PRJNA1255621 | Arabidopsis RNA | Control | Control-4 | 23435544 | 95.66% |
| SAMN48150801 | PRJNA1255621 | Arabidopsis RNA | Lead | Lead-1 | 21166326 | 96.59% |
| SAMN48150802 | PRJNA1255621 | Arabidopsis RNA | Lead | Lead-2 | 25742964 | 96.56% |
| SAMN48150803 | PRJNA1255621 | Arabidopsis RNA | Lead | Lead-3 | 26789286 | 96.32% |
| SAMN48150804 | PRJNA1255621 | Arabidopsis RNA | Lead | Lead-4 | 31105096 | 96.84% |
